# Scalable Production of a *De Novo* SARS-CoV-2 Antiviral miniprotein in E*scherichia coli*

**DOI:** 10.64898/2026.06.23.734092

**Authors:** Jinhwan Shin, Eu-min Kim, Jun-hong Jang, Seok-won Jee, Sang-hoon Kim, Seonggwan Yu, Minguen Yoon, Daniel Craig, Ryan Swoyer, Sandip Patel, Praveen Alamuri, Albert Price, Rashmi Ravichandran, Lauren Carter, Sammaiah Pallerla

## Abstract

The rapid emergence of SARS-CoV-2 variants that evade neutralizing antibodies underscores the need for next-generation antiviral biologics that combine molecular precision with scalable, cost-effective manufacturing. Computationally designed miniproteins targeting the receptor-binding domain (RBD) of the spike protein offer a compelling alternative to monoclonal antibodies due to their small size, high thermal stability, and compatibility with microbial expression systems. Here we report the end-to-end development and cGMP production of IPD-52520, a *de novo* antiviral miniprotein, using an optimized *E. coli* platform. Two miniprotein candidates, a homotrimeric construct (Trimer is referred to as IPD-52520, 17 kDa) and a tandem fusion (Daisy is referred to as IPD-52521, 25 kDa), were evaluated in parallel through systematic optimization of strain selection, media composition, fed-batch fermentation, inclusion-body solubilization, refolding, and chromatographic purification. The Trimer was downselected as the lead molecule based on superior preclinical efficacy, favorable pharmacokinetic properties, and higher volumetric manufacturing yields. The optimized process delivers approximately 2 g/L of purified protein at greater than 90% purity. Scale-up from 5 L to 50 L under cGMP conditions demonstrated excellent batch-to-batch reproducibility across six independent batches, supporting nonclinical and Phase 1 clinical supply. Comprehensive biophysical characterization confirmed a well-folded, predominantly alpha-helical trimer (*T_m_* = 73.4 °C; polydispersity = 1.005) with an intact primary structure and strong target-binding affinity (*K_D_* < 1 pM). Real-time stability studies indicate that the drug substance is stable at 2–8 °C for at least 12 months, with ongoing stability studies. These results demonstrate the feasibility of translating computationally designed antiviral miniproteins into manufacturable biologics and provide a platform applicable to rapid-response therapeutics against current and future pandemic threats.

## 1. Introduction

The COVID-19 pandemic exposed a critical gap in global preparedness: the inability to rapidly design, produce, and deploy effective antiviral biologics at scale. Ongoing evolution of SARS-CoV-2 has eroded the efficacy of neutralizing antibodies elicited by infection or vaccination, rendering many monoclonal antibody therapies ineffective against emerging variants [1]. These challenges underscore the urgent need for next-generation antivirals that combine molecular precision with rapid, scalable, and cost-effective manufacturing.

The convergence of artificial intelligence, computational protein design, and high-throughput experimental validation has transformed our ability to engineer novel proteins from first principles [2–4]. Advances in structure-prediction algorithms such as AlphaFold and RoseTTAFold, combined with generative design methods including RFdiffusion and ProteinMPNN, now enable the creation of compact, well-defined proteins with tailored binding properties that far exceed natural capabilities [5–9].

Miniproteins; small, structured proteins typically comprising 40–80 amino acids, represent a particularly promising therapeutic modality that bridges the gap between monoclonal antibodies and small-molecule drugs [10–12]. These *de novo*-designed molecules offer several inherent advantages: small size enables enhanced tissue penetration; exceptional thermal stability eliminates cold-chain requirements; compatibility with microbial expression systems reduces manufacturing complexity; and resistance to proteolytic degradation [13,14]. Unlike traditional peptide therapeutics, which often suffer from poor stability and rapid clearance, miniproteins can be engineered to achieve antibody-like binding affinities while maintaining the pharmacokinetic advantages of smaller molecules [12].

The Institute for Protein Design (IPD) at the University of Washington has pioneered *de novo* miniprotein therapeutics, demonstrating their potential across multiple therapeutic areas [15]. Foundational work by Chevalier *et al*. established massively parallel design-build-test cycles, generating more than 22,000 miniproteins that target influenza hemagglutinin and botulinum neurotoxin with picomolar binding affinities [18]. Subsequent advances enabled the design of miniproteins against previously ‘undruggable’ targets, including protein-protein interfaces and conformationally dynamic targets [17,18].

SARS-CoV-2 served as an ideal testbed for computational miniprotein design. In landmark work, Cao *et al*. reported the *de novo* design of picomolar SARS-CoV-2 miniprotein inhibitors targeting the RBD of the spike protein, demonstrating neutralization activity rivaling that of the best-known monoclonal antibodies [19]. These designed molecules, including the lead candidates LCB1 and LCB3, exhibited exceptional stability, retaining activity after prolonged exposure to elevated temperatures. Hunt *et al*. subsequently developed multivalent architectures that enhanced avidity-driven binding, while Case et al. demonstrated in vivo efficacy, providing prophylactic and therapeutic protection in mouse models [20,21].

Beyond SARS-CoV-2, the miniprotein platform has expanded to address a range of therapeutic challenges. Recent applications include miniprotein antagonists for autoimmune diseases that target IL-23R and IL-17A and have oral bioavailability [22], ultrahigh-affinity PD-L1 binders for cancer immunotherapy and molecular imaging [23], and engineered miniproteins for bacterial toxin neutralization [24]. The first computationally designed protein medicine, SKYCovione, a COVID-19 vaccine developed in collaboration with SK Bioscience, has received regulatory approval, validating the translational potential of this approach [25].

Despite these advances, a critical gap remains between computational design and manufacturable therapeutics. Although miniprotein design algorithms can generate high-affinity binders [26], the development of robust, scalable bioprocesses for their production has received limited attention. Recombinant protein production in *Escherichia coli* is an attractive platform for non-glycosylated miniproteins, offering established infrastructure, cost efficiency, and regulatory precedent. However, expressing structurally complex *de novo* proteins at high yield remains challenging due to folding difficulties, inclusion body formation, and aggregation during high-level bacterial expression.

Here, we report the development, cGMP scale-up, and comprehensive characterization of IPD-52520, a novel antiviral miniprotein targeting SARS-CoV-2, produced in an optimized *E. coli* expression system. We establish an integrated framework spanning expression optimization, upstream and downstream process development, protein characterization, and stability evaluation. This work demonstrates the feasibility of translating computationally designed antivirals into manufacturable biologics and provides a platform for rapid-response therapeutics against current and future pandemic threats.

## 2. Materials and Methods

### 2.1 Plasmid Preparation and Confirmation

Two miniprotein candidates, Trimer (IPD-52520) and Daisy (IPD-52521), were obtained from the Institute for Protein Design (IPD) and were commercially synthesized by GenScript. Plasmids were provided with Certificates of Analysis and verified by sequencing. The Trimer plasmid matched the reference (C389-AHB2v1-2GS-SB175_tagless_pET29b); the Daisy plasmid contained four nucleotide differences relative to the reference (AHB2v2_12PAS_LCB3v2.2_12PAS_LCB1v2.2_pET29b), which were confirmed as silent mutations with no changes to the encoded amino acid sequence.

### 2.2 Bacterial Strain Selection

Six *E. coli* strains were screened for miniprotein expression: BL21(DE3) (Enzynomics), Rosetta(DE3), Rosetta-gami2(DE3), Tuner(DE3) (Novagen), C41(DE3), and C43(DE3) (Lucigen). Initial screening was conducted in shake-flask cultures, followed by 5 L fed-batch fermentation of the three highest-expressing strains. Protein expression was assessed by SDS-PAGE using 20-fold-diluted cell-pellet samples.

### 2.3 Cell Bank System

#### Pre-Master Cell Bank

The pre-MCB was produced in SK Bioscience’s R&D facility. The purified expression plasmid (52520, C389-AHB2v1-2GS-SB175_pET-29b(+)_tagless) was extracted from IPD-supplied cells (MB-RWCB-52520) using an AccuPrep Nano-Plus Plasmid Mini Extraction kit and heat-shock transformed into *E. coli* BL21(DE3) (One Shot, Thermo Fisher, Cat. No. 44-0048). Transformants were recovered in 1 mL of LB at 37 °C and 200 rpm for 1 h, plated on LB+kanamycin agar, and incubated overnight. A single colony was expanded through four serial passages in 128 mL of soytone–yeast extract (SY) medium supplemented with glucose, kanamycin, MgCl₂, and trace metals until no lag phase was observed, confirming adaptation. Culture purity was verified by Gram staining and microscopy. Glycerol stocks (20% v/v) were stored at ≤ −70 °C in 1 mL aliquots. Cell viability and plasmid retention were determined by serial dilution plating on LB and LB–kanamycin agar.

#### Master and Working Cell Banks

The MCB and WCB were produced by SK Bioscience (Andong, South Korea) from pre-MCB glycerol stocks. A 37.5 µL inoculum from a thawed pre-MCB vial was expanded through two passages in SY medium (OD₆₀₀ > 4.0 in each passage). After Gram-stain confirmation of purity, glycerol stocks (20% v/v) were aliquoted and stored at ≤ −135 °C (MCB, 100 vials; WCB, 200 vials). Plasmid copy number was determined by sequence-specific quantitative PCR (qPCR) (128 copies per genome equivalent). The cell bank testing program is summarized in Table 1.

**Table 1.** Cell bank testing program for MCB and WCB.

| Test | MCB | WCB |
| --- | --- | --- |
| Cell viability | Pass | Pass |
| Plasmid retention | Pass | Pass |
| Gram stain / Morphology | Gram-negative rods | Gram-negative rods |
| Plasmid copy number (qPCR) | 128 copies/genome eq. | N/A |
| Sterility / Contamination | No contamination | No contamination |

### 2.4 Fermentation Process Development

#### Media Optimization

Two media were compared: RRTBII (IPD reference medium) and SY medium (SK Bioscience; 20 g/L Difco Soytone, 10 g/L Bacto Yeast Extract, 6 g/L NaCl). Fed-batch fermentation was conducted in 5 L bioreactors at 37 °C ± 1 °C, pH 6.95 ± 0.05, and 30% dissolved oxygen.

#### Feed Medium and Feeding Strategy

A control feed medium (200 g/L glucose, 100 g/L yeast extract, 18.75 mM MgClLJ) was benchmarked against a high-glucose formulation (500 g/L). Two feeding methods were compared: manual pump control and pH-stat feeding, in which the feed medium replaced acid addition to maintain pH as the carbon source was consumed.

#### Process Parameter Optimization

Initial glucose concentrations of 1, 10, and 30 g/L were tested. Aeration was optimized at gas flow rates of 2.0, 4.0, and 6.0 SLPM. Cell growth was monitored by OD₆₀₀.

#### Induction Optimization

Expression was induced with Isopropyl β-D-1-thiogalactopyranoside (IPTG). Three parameters were optimized: induction timing (early vs. late, based on ODLJLJLJ), IPTG concentration (5, 7, and 11 mL of 0.5 M IPTG per 2.5 L culture), and induction duration (samples collected every 2 h post-induction). All samples were analyzed by SDS-PAGE.

### 2.5 Recovery and Purification Development

#### Cell Lysis and Solubilization

Four lysis methods were evaluated: heat lysis (70 °C), microfluidization (850–1000 bar), osmolysis (osmotic shock), and guanidine lysis. For guanidine solubilization, chaotropic agents (guanidine-HCl vs. urea) were compared at concentrations of 1–6 M, and buffer volume was optimized at 3X, 5X, and 10X the cell pellet weight. The optimized solubilization buffer contained 6 M guanidine-HCl in 1X Dulbecco’s Phosphate Buffered Saline (DPBS) with 0.05% Polyethyleneimine (PEI) for endotoxin removal (room temperature, 4 h, gentle mixing).

#### Refolding

Two refolding methods were compared: dilution refolding (1:9) and TFF diafiltration (≥4 diavolumes, 1X DPBS, pH 7.4 ± 0.2, TMP ≤ 1.0 bar, 15–25 °C, ≤ 24 h).

#### Chromatographic Purification

Multiple resins were screened, including AEX (DEAE-FF, Capto DEAE, Sartobind Q, Q-FF, ANX-FF), CEX (SP-FF), mixed-mode (hydroxyapatite), and HIC (Phenyl-FF). DEAE-FF chromatography was performed in 1x DPBS (pH 7.4 ± 0.2) with elution using a linear NaCl gradient (0–40% of 1 M NaCl in DPBS). Dynamic binding capacity, gradient mode (linear vs. step), and secondary polishing strategies were evaluated.

#### UF/DF and Endotoxin Removal

UF/DF was performed using 30 kDa MWCO membranes (≥10 diavolumes, TMP ≤1.0 bar, 15–25 °C). Endotoxin removal combined with PEI treatment (0.01–0.1%) during solubilization with a secondary Pierce High-Capacity Endotoxin Removal Resin step. Endotoxin was quantified by kinetic chromogenic LAL assay (EU/µg protein).

### 2.6 Analytical Methods

Cell growth was monitored by OD₆₀₀. Culture purity was confirmed by Gram staining. Protein concentration was determined by Bradford assay and UV₂₈₀. Purity and molecular weight were assessed by SDS-PAGE (gradient gels) and SEC-HPLC. Target molecular weights were 17 kDa (Trimer) and 25 kDa (Daisy). Cell harvest was performed by centrifugation at 12,000 x g for 15 min.

## 3. Results

### 3.1 Strain Selection

Flask-scale screening identified BL21(DE3), C41(DE3), and C43(DE3) as the highest-expressing strains by SDS-PAGE (Figure 1). Subsequent 5 L fed-batch fermentation confirmed BL21(DE3) as optimal, with superior expression of both Trimer and Daisy compared with C41(DE3) and C43(DE3) (Figure 2 and 3).

**Figure 1.**
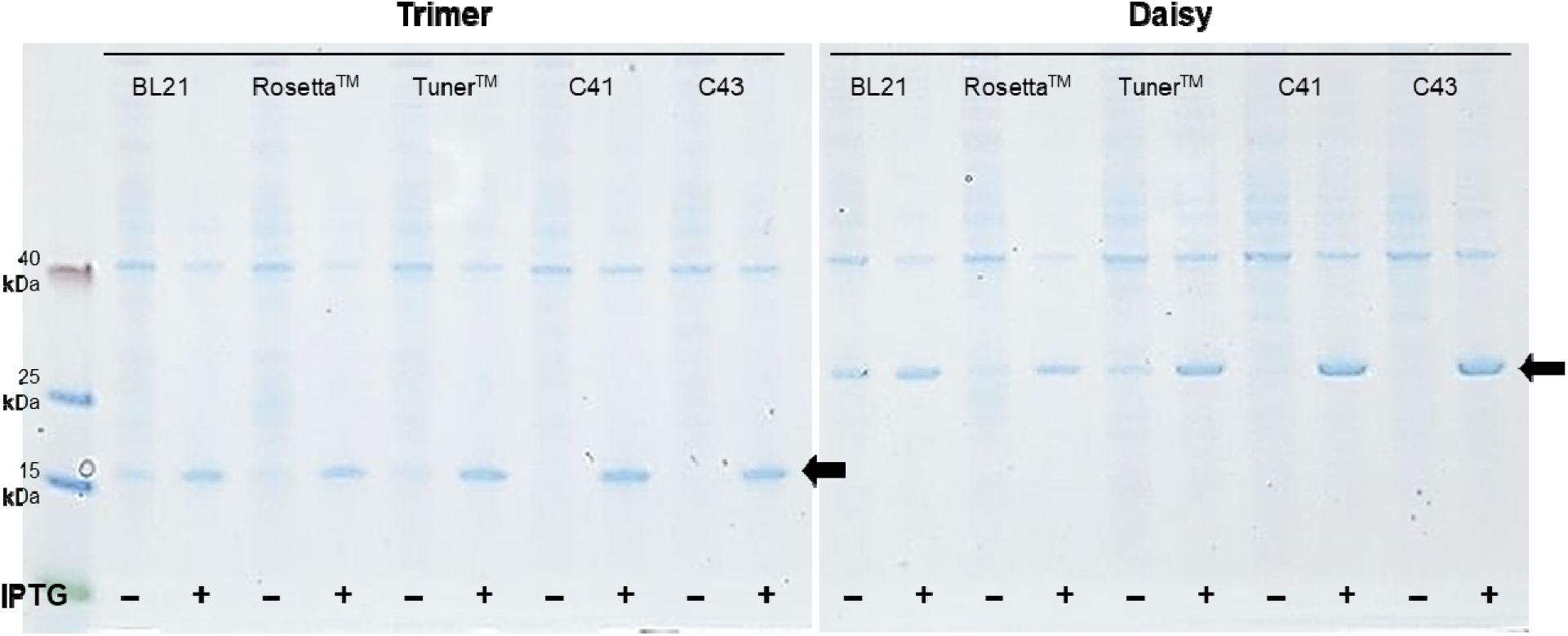
Miniprotein expression in six *E. coli* strains. Cell pellets were diluted 20-fold and analyzed by SDS-PAGE. (−) pre-induction; (+) post-induction. Arrows indicate Trimer and Daisy bands. BL21(DE3), C41(DE3), and C43(DE3) showed the highest expression. Rosetta-gami2(DE3) displayed low target expression under the tested induction conditions and was therefore not advanced to the comparative SDS-PAGE shown in Figure 1.

**Figure 2.**
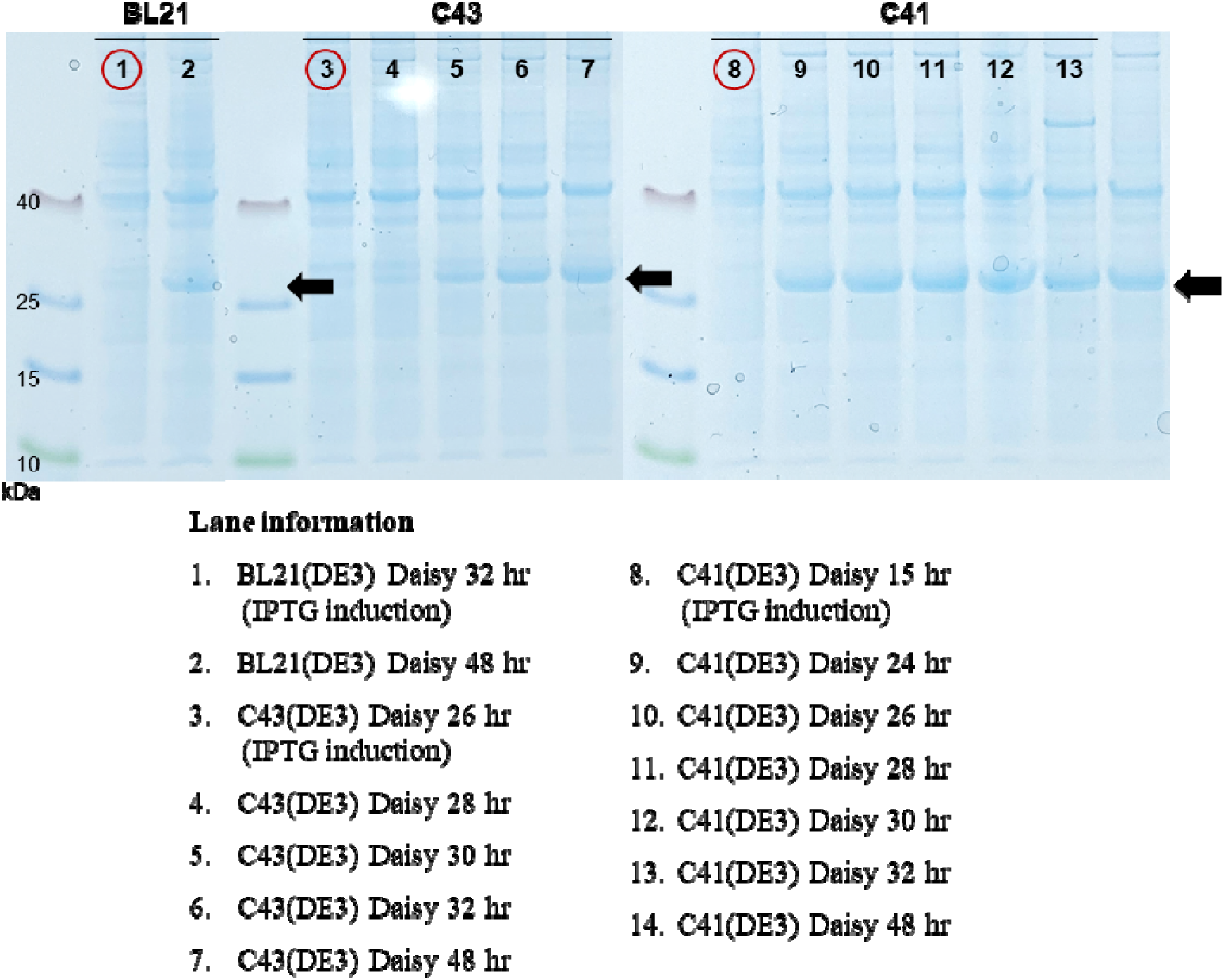
Daisy expression in BL21(DE3), C43(DE3), and C41(DE3) during 5 L fed-batch fermentation. Red circles, IPTG induction; black arrows, miniprotein bands. BL21(DE3) produced the highest yields.

**Figure 3.**
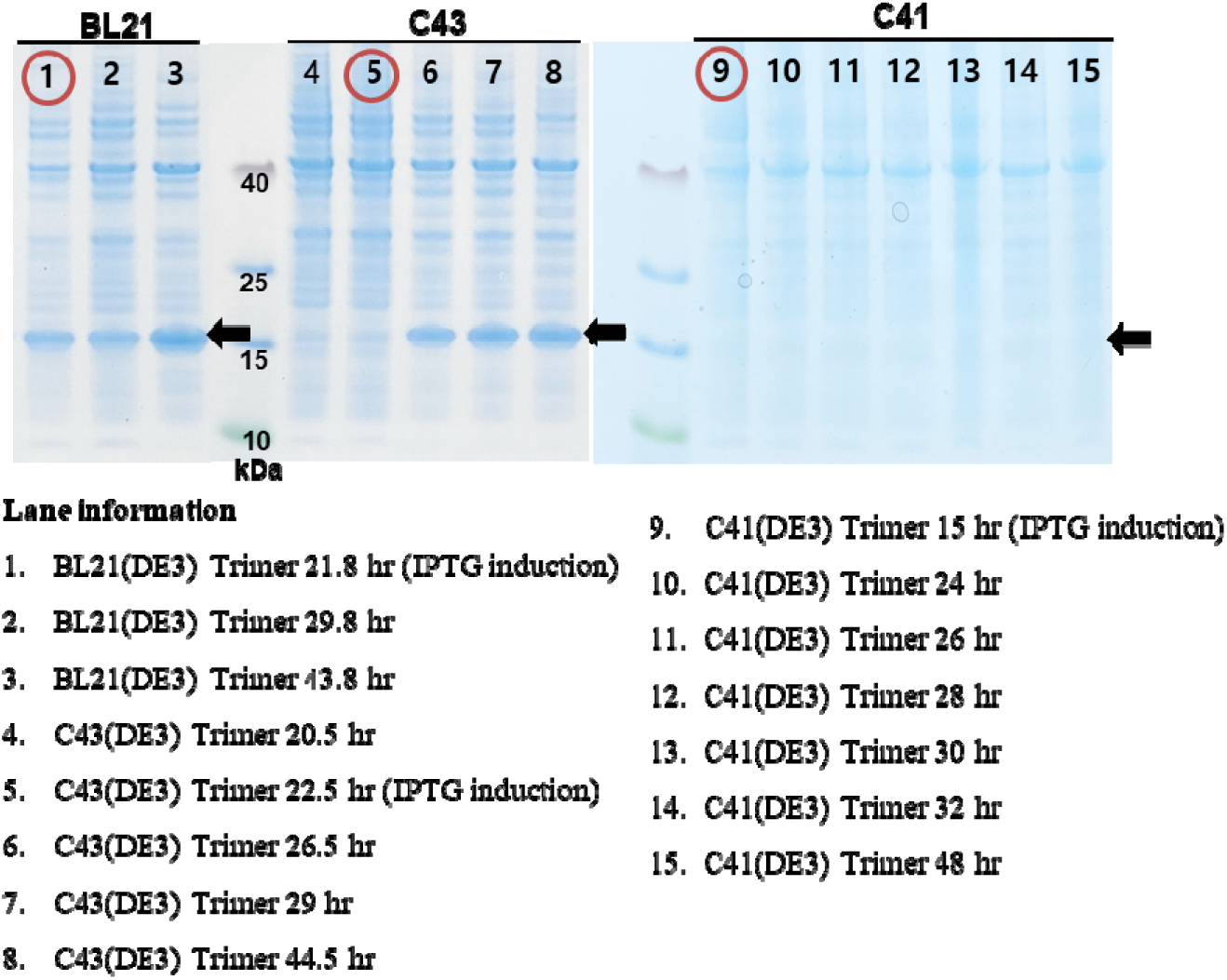
Trimer expression in BL21(DE3), C43(DE3), and C41(DE3) during 5 L fed-batch fermentation. Red circles, IPTG induction; black arrows, miniprotein bands. BL21(DE3) produced the highest yields.

### 3.2 Media and Feeding Strategy Optimization

SY medium outperformed RRTBII in several respects (Table 2; Figure 4): earlier induction (26.0 vs. 33.0 h), comparable final cell density (OD₆₀₀ 82.3 vs. 80.6), and similar cell pellet yield (793.1 vs. 776.7 g). Its simpler three-component formulation offers clear advantages for industrial-scale production.

**Figure 4.**
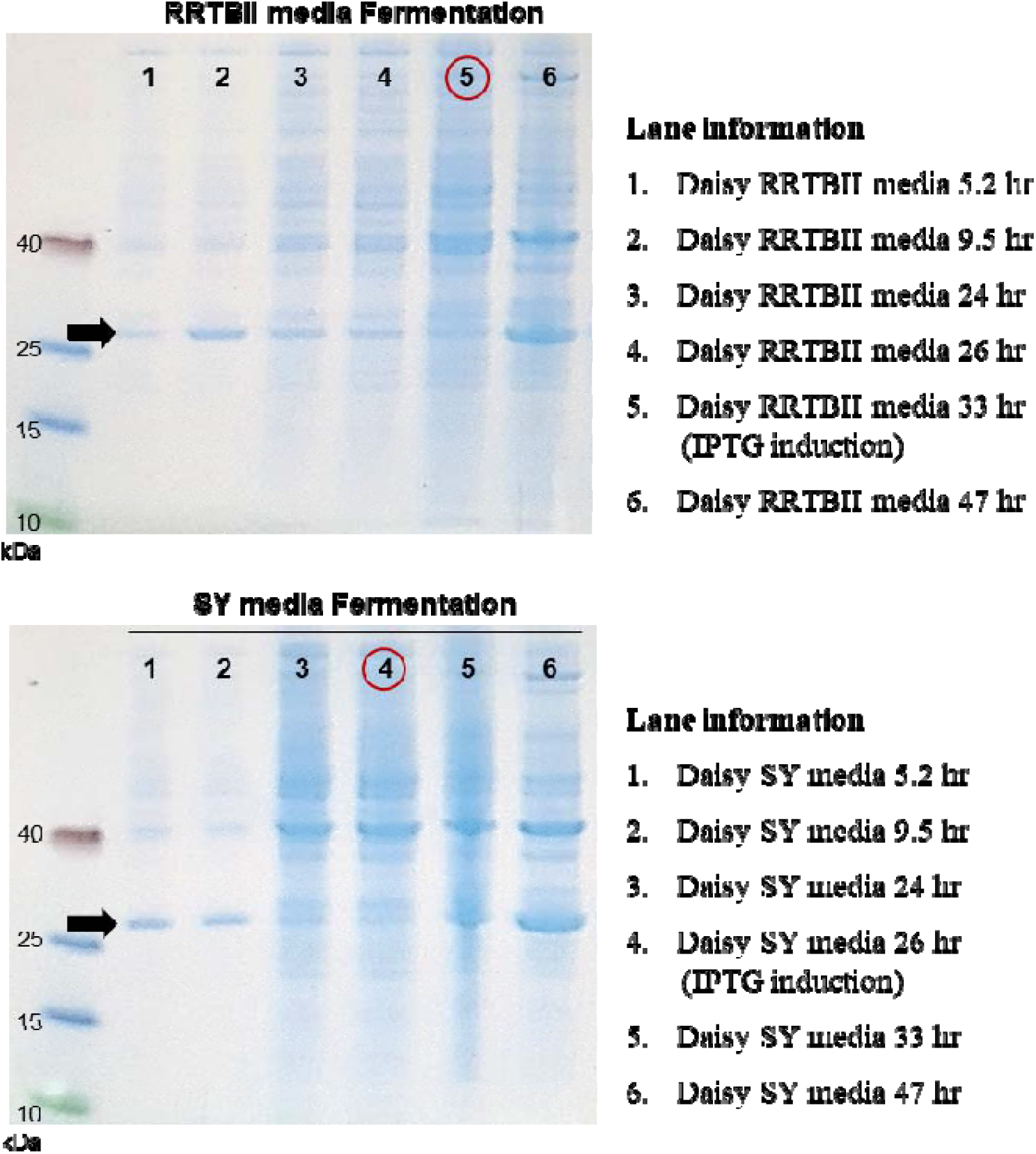
Daisy expression in RRTBII vs. SY media. Red circles, IPTG induction; black arrows, Daisy bands. SY medium enabled earlier induction and higher expression.

**Table 2.** Fermentation performance: RRTBII vs. SY medium.

| Parameter | RRTBII | SY |
| --- | --- | --- |
| Induction time (h) | 33.0 | 26.0 |
| Final OD<sub>600</sub> | 80.6 | 82.3 |
| Cell pellet yield (g) | 776.7 | 793.1 |

The standard feed medium (200 g/L glucose) outperformed the 500 g/L formulation, which inhibited growth and protein expression. pH-stat feeding surpassed manual pump control, yielding higher final cell density (OD₆₀₀ 105.0 vs. 88.1) and greater cell-pellet yield (910.8 vs. 578.8 g, a 57% increase).

### 3.3 Process Parameter and Expression Optimization

An initial glucose concentration of 10 g/L provided the best balance of growth kinetics and process efficiency. Aeration at 4.0 SLPM optimized the trade-off between growth rate and expression yield. Late induction (OD₆₀₀ ∼74) significantly outperformed early induction (OD₆₀₀ ∼4), yielding higher final cell density and improved protein expression (Figure 5). The original IPTG concentration (1 mM) was optimal, and 6–8 h post-induction was sufficient to achieve maximum expression. Optimized parameters are summarized in Table 3.

**Figure 5.**
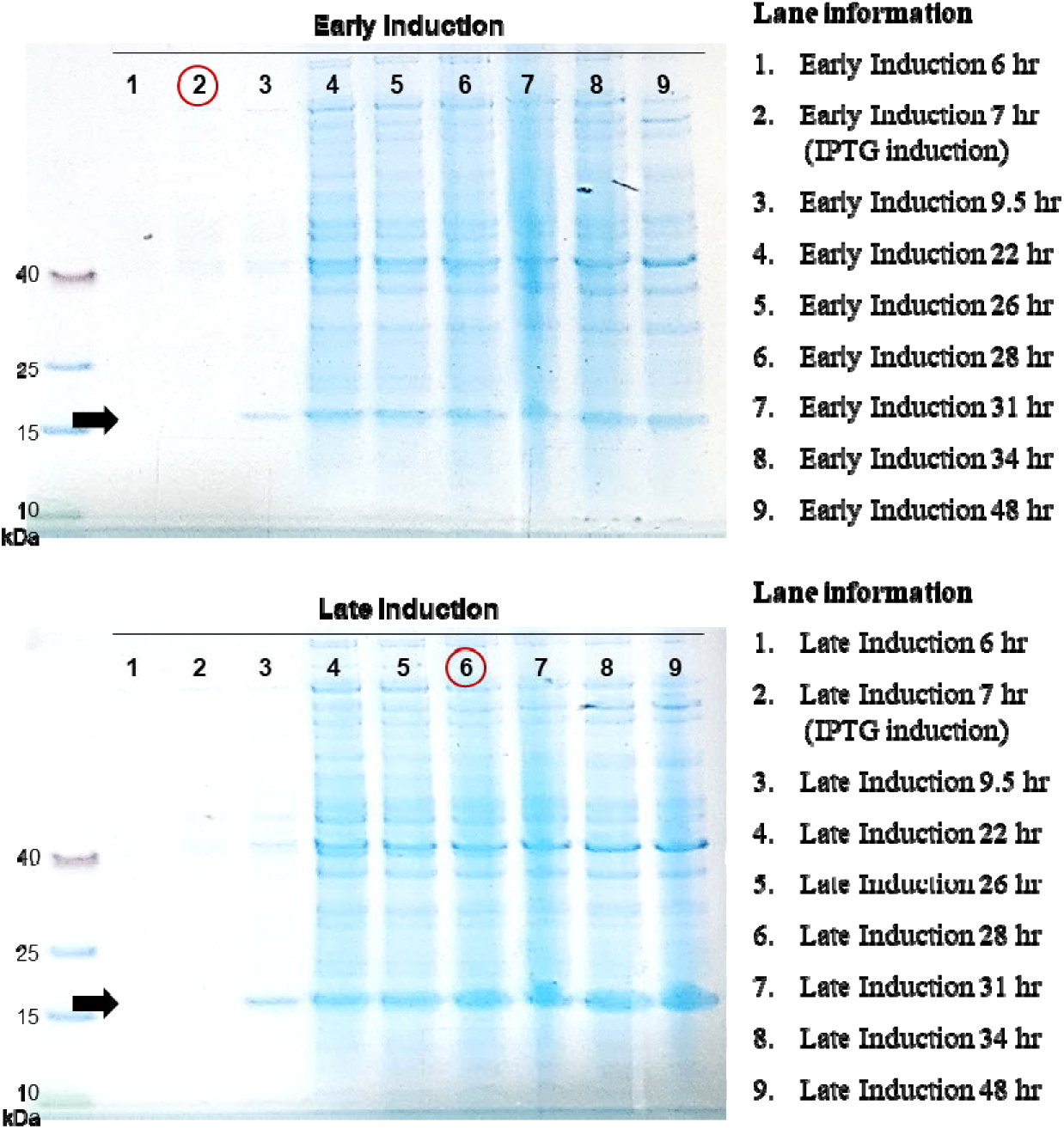
Trimer expression comparing early vs. late induction in 5 L fermentation. Red circles, IPTG induction; black arrows, miniprotein bands. Late induction achieved substantially higher expression.

**Table 3.** Optimized fermentation parameters for SARS-CoV-2 miniprotein production.

| Parameter | Optimized Condition |
| --- | --- |
| *E. coli* strain | BL21(DE3) |
| Culture medium | SY medium |
| Initial glucose | 10 g/L |
| Feeding strategy | pH-stat |
| Gas flow rate | 4.0 SLPM |
| Induction timing | OD<sub>600</sub> ≥60 (late) |
| IPTG concentration | 1 mM |
| Induction duration | 6–8 h |

### 3.4 Recovery Method Optimization

Comparative evaluation of lysis methods revealed marked differences in yield and purity (Table 4). Heat lysis achieved high purity (≥90.9%) but extremely low yields (≤91 mg/L for Trimer). Microfluidizer lysis improved yields but reduced purity. Osmolysis produced variable results—high Daisy yield (1,881 mg/L) but poor Trimer recovery (306 mg/L).

**Table 4.** Lysis method comparison for miniprotein recovery (after DEAE-FF chromatography).

| Lysis Method | Protein | Yield (mg/L) | Purity (%) |
| --- | --- | --- | --- |
| Heat lysis | Daisy | 145 | 99.3 |
|  | Trimer | 91 | 90.9 |
| Microfluidizer | Daisy | 669 | 82.0 |
|  | Trimer | 585 | 80.5 |
| Osmolysis | Daisy | 1,881 | 90.0 |
|  | Trimer | 306 | 97.2 |

Guanidine-HCl was selected for inclusion-body solubilization. It proved superior to urea, achieving complete solubilization of the Trimer at 6 M (Figure 6). A 5X buffer volume (relative to cell-pellet weight) maintained yield while reducing process volume.

**Figure 6.**
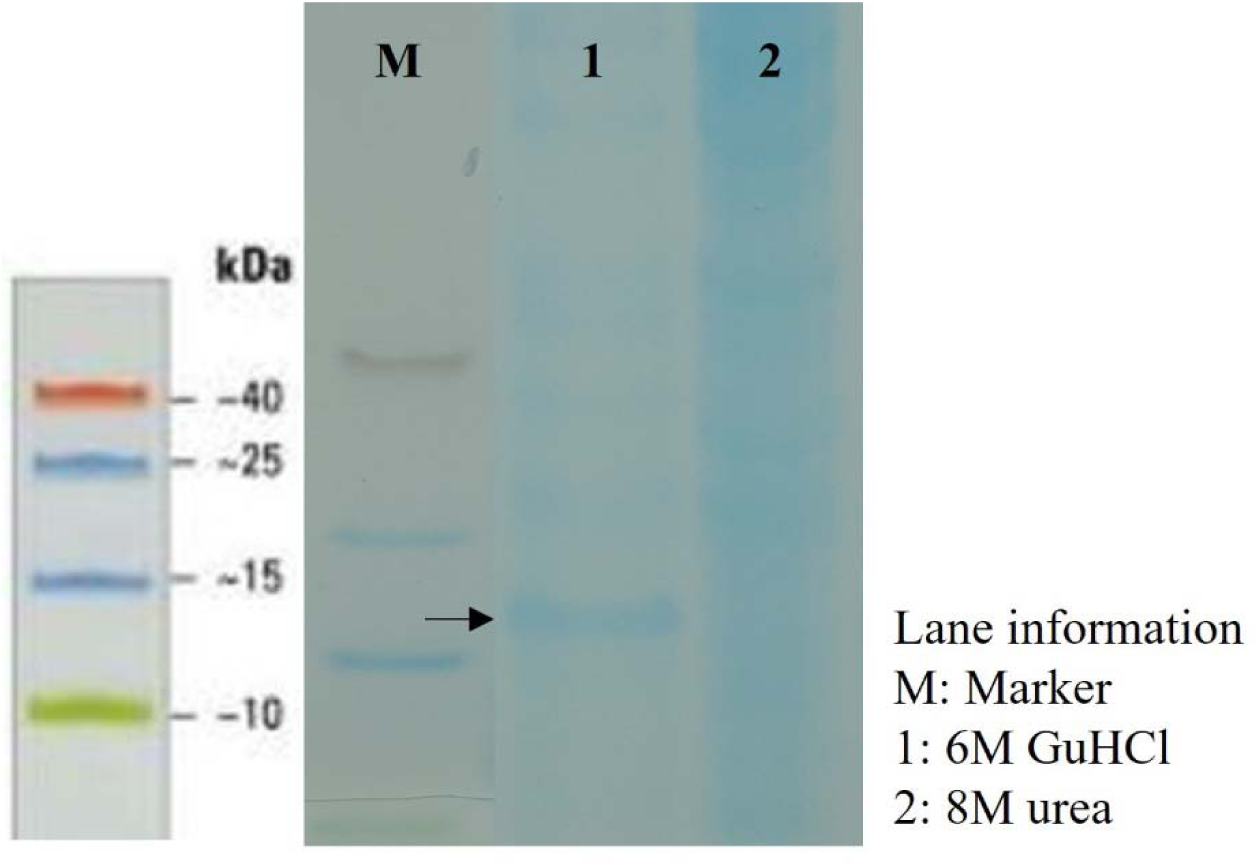
SDS-PAGE comparison of chaotropic agents. Guanidine-HCl achieved complete Trimer solubilization; urea did not.

**Figure 7.**
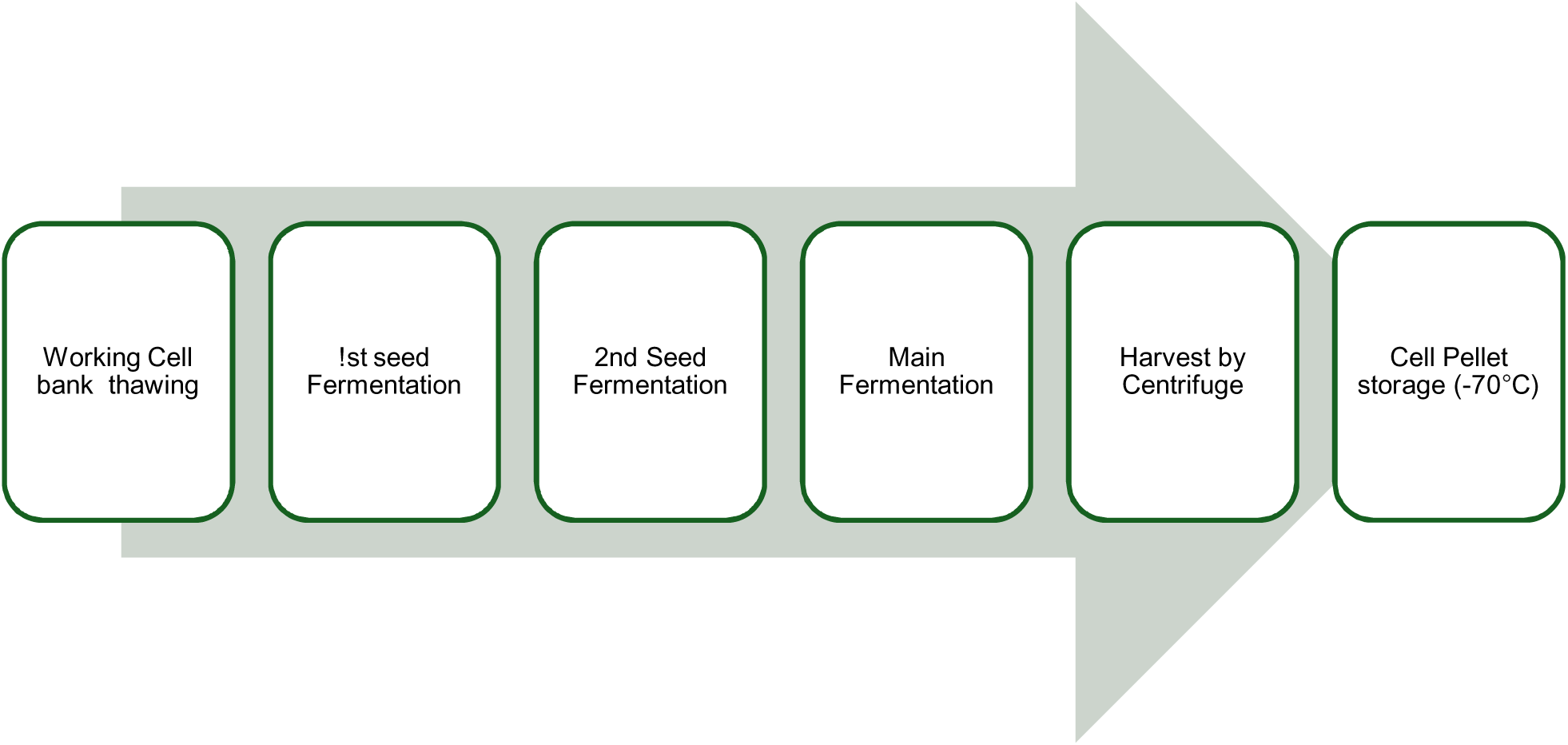
Process flow Schema for 50 L cGMP upstream manufacturing of IPD-52520.

### 3.5 Refolding and Chromatographic Purification

Tangential Flow Filtration (TFF) diafiltration outperformed dilution refolding by enabling complete guanidine removal at constant volume with substantially lower buffer consumption.

Among AEX resins, DEAE-FF was superior (Table 5): it delivered a higher step yield (50.5% vs. 46.2% for Capto DEAE) and better impurity removal. Sartobind Q failed to bind the target protein. Secondary screening confirmed that CEX (SP-FF), mixed-mode (hydroxyapatite), and alternative AEX resins (Q-FF, ANX-FF) could not effectively resolve High Molecular Weight (HMW) impurities. Linear gradient elution with DEAE-FF (DBC: 50 mg/mL resin) provided optimal separation. UF/DF with 30 kDa MWCO membranes increased purity from approximately 75–85% to >90%, primarily by removing Low-Molecular-Weight (LMW) species.

**Table 5.**
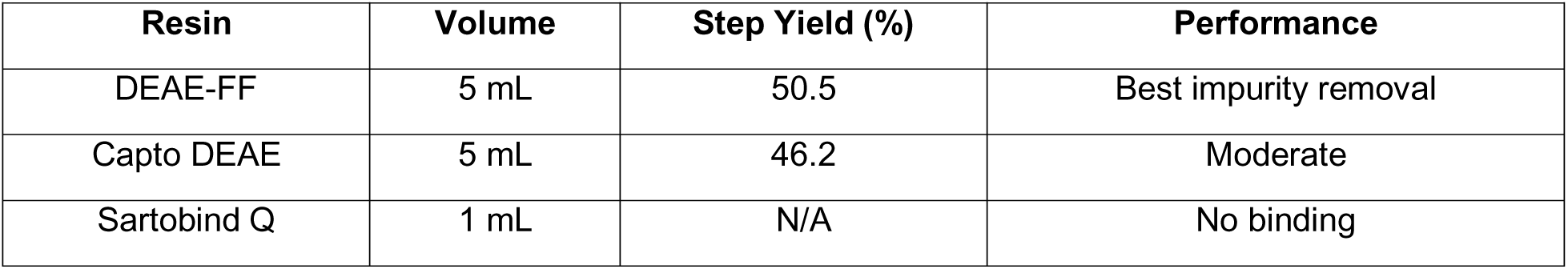
Anion exchange resin screening results.

| Resin | Volume | Step Yield (%) | Performance |
| --- | --- | --- | --- |
| DEAE-FF | 5 mL | 50.5 | Best impurity removal |
| Capto DEAE | 5 mL | 46.2 | Moderate |
| Sartobind Q | 1 mL | N/A | No binding |

### 3.6 Endotoxin Removal

PEI at 0.05% achieved effective endotoxin reduction while preserving protein recovery; concentrations ≥0.1% caused approximately 50% yield loss. Subsequent treatment with an endotoxin-removal resin reduced levels from 0.16–0.21 EU/µg to 0.02–0.03 EU/µg (86–88% removal efficiency).

### 3.7 Scale-Up Confirmation

During the initial development stage, scale-up from 1 L to 10 L showed excellent reproducibility (Table 6). Daisy yields were 1.7– 2.0 mg/L at 94.4–98.7% purity; Trimer reached 2.5 -2.9 mg/L at 93.1–95.8% purity.

**Table 6.** Scale-up validation: laboratory vs. pilot-scale purification.

| Protein | Scale | Yield (mg/L) | Purity (%) |
| --- | --- | --- | --- |
| Daisy | Lab (1 L) | 1,956 | 98.7 |
|  | Pilot (10 L) | 1,733 | 94.4 |
| Trimer | Lab (1 L) | 2,490 | 95.8 |
|  | Pilot (10 L) | 2,925 | 93.1 |

## 4. Discussion

### 4.1 Fermentation Process

Choosing BL21(DE3) is consistent with its well-established effectiveness in producing recombinant proteins, thanks to genetic modifications that enhance expression and reduce proteolysis. Switching from RRTBII to SY medium reduced process time, simplified the formulation from 6 to 3 components, and maintained high expression levels—benefits that lead to lower raw material costs, easier quality control, and less regulatory burden at industrial scale.

The pH-stat feeding strategy automatically controls nutrient delivery by exploiting the natural pH decrease during carbon-source consumption. This approach resulted in a 57% increase in cell-pellet mass compared to manual pump control. Setting the initial glucose concentration at 10 g/L balanced quick early growth with the risk of catabolite repression. An aeration rate of 4.0 SLPM ensured sufficient oxygen transfer without causing excessive shear stress. Inducing at late stages (OD₆₀₀ ≥60) maximized the cellular machinery for protein production. A 6-8 hour induction period provided sufficient time for accumulation while reducing degradation and the formation of inclusion bodies.

### 4.2 Recovery and Purification

The poor performance of soluble-protein recovery methods confirmed that both miniproteins predominantly form inclusion bodies in *E. coli*, prompting a shift to guanidine-based solubilization. Guanidine-HCl outperformed urea due to its stronger chaotropic activity. TFF-based refolding provided superior process control, complete denaturant removal, and dramatically reduced buffer consumption compared with dilution refolding.

DEAE-FF emerged as the optimal chromatographic resin, with high dynamic binding capacity (50 mg/mL), good step yields (>50%), and effective impurity clearance. The inability of multiple orthogonal chromatographic modes (CEX, HIC, MMC, and alternative AEX) to resolve HMW species suggests that these contaminants share physicochemical properties with the target proteins and may represent partially aggregated or misfolded product forms rather than host-cell proteins. Protein aggregation near the isoelectric points (pI 4.63/4.71) during CEX supports this interpretation. UF/DF with 30 kDa MWCO membranes proved essential for LMW impurity removal, consistently increasing purity from 75-85% to >90%.

#### 4.2.1 Evolution of the Downstream Process: Introduction of HIC, Secondary PEI Addition, and AEX Detergent Wash

The preliminary downstream process, developed using material from 5 L and 10 L fermentation runs, provided the foundation for cGMP manufacturing but required further optimization before it could reliably meet the proposed drug substance endotoxin specification at the 50 L scale. The primary challenge was achieving consistent, adequate endotoxin clearance without incurring unacceptable yield losses.

A systematic laboratory-scale development study was conducted to evaluate several independent process modifications: increased PEI concentration in the solubilization buffer, optimization of solubilization container geometry, increased post-solubilization centrifuge speed, introduction of post-refolding depth filtration, replacement of the stand-alone endotoxin removal resin step with a hydrophobic interaction chromatography (HIC) polishing step using Phenyl Sepharose Fast Flow resin followed by relocation of the second UF/DF step, introduction of a second PEI addition step after refolding, and a detergent wash of the AEX column using Tergitol prior to product elution.

Of the approaches evaluated, three-unit operations provided the best balance between endotoxin reduction and retention of process yield and were advanced to 50 L confirmation runs: (i) introduction of HIC polishing, (ii) a second post-refolding PEI addition step, and (iii) a Tergitol detergent wash of the AEX resin. The rationale for each merits a brief discussion. The HIC step exploits the relatively hydrophobic character of endotoxin acyl chains, enabling selective retardation of lipopolysaccharide under moderate ammonium sulfate conditions while the IPD-52520 elutes earlier; this orthogonal separation mode complements the charge-based selectivity of AEX and addresses the HMW impurity profile discussed above. The second PEI addition step targets residual endotoxin in the refolded pool prior to chromatographic capture, using the well-established mechanism of PEI–endotoxin electrostatic complexation and co-precipitation, at a concentration (0.03% v/v) determined to minimize co-precipitation of the target protein. The Tergitol (polyethylene glycol tert-octylphenyl ether) wash of the AEX column disrupts hydrophobic interactions between endotoxin and the resin that can persist through standard salt washes, releasing endotoxin prior to product elution and thereby reducing the endotoxin burden in the AEX eluate pool.

The first 50 L confirmation run, conducted with 1% Tergitol in the AEX column wash step, produced material that did not meet the proposed drug substance endotoxin specification. Analysis of in-process samples indicated that a 1% Tergitol concentration was insufficient to fully displace the hydrophobic endotoxin fraction adsorbed to the DEAE resin under the process loading and wash conditions employed. In response, the Tergitol concentration was increased to 3% in the second 50 L confirmation run. This modification produced drug substance that met all proposed specifications, including the endotoxin limit, without significant impact on step yield or purity, confirming that 3% Tergitol represents an effective and process-compatible endotoxin clearance measure for this resin and load condition. The 3% Tergitol AEX wash was therefore incorporated into the cGMP process and is represented accordingly in the downstream process flow diagram (Figure 8). Together, these three-unit operations- HIC polishing, secondary PEI precipitation, and AEX detergent wash- constitute a layered, orthogonal endotoxin clearance strategy that reflects current best practices for the purification of *E. coli* inclusion body-derived therapeutics.

**Figure 8.**
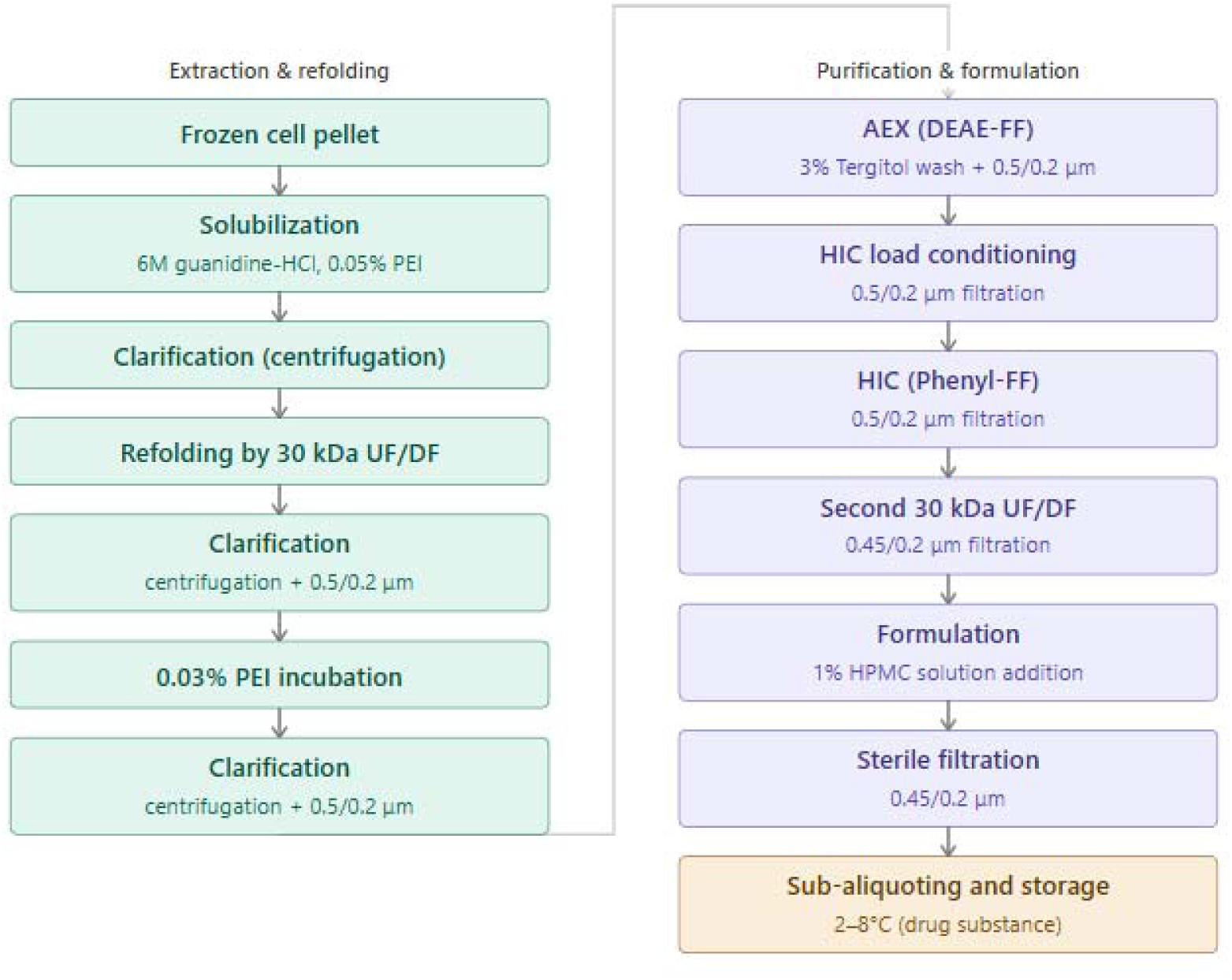
Process flow diagram for cGMP downstream purification of IPD-52520.

#### 4.2.2 Second UF/DF Step, Formulation, and Clarification of HPMC Concentration

Following HIC chromatography, a second ultrafiltration/diafiltration step (UF/DF2) was performed using 30 kDa nominal molecular weight cutoff membranes to serve two essential, sequential purposes: removal of ammonium sulfate carried over from the HIC elution pool and concentration of IPD-52520 to a protein level suitable for final formulation. The HIC eluate pool, which contains approximately 1 M ammonium sulfate due to the descending salt gradient used to elute the target protein, is incompatible with the drug substance formulation buffer and must be fully exchanged prior to HPMC addition. Following concentration to a retentate mass of approximately 0.5 kg, the material underwent a minimum of ten diavolumes against DPBS pH 7.2 as the diafiltration buffer, with diafiltration endpoint defined by an inline permeate conductivity of 11–17 mS/cm. Upon completion of diafiltration, the retentate and the DPBS rig flush were combined, and the protein concentration was measured by absorbance at 280 nm. A calculated volume of the flush was then added to the retentate to achieve a target IPD-52520 concentration of 66.0 g/L. The UF/DF2 pool was subsequently clarified by 0.45/0.2 µm filtration prior to formulation.

The clarified UF/DF2 filtrate was formulated by adding hydroxypropyl methylcellulose (HPMC), a non-ionic cellulose polymer selected as a stabilizing excipient to reduce surface adsorption and mitigate aggregation at the high protein concentrations required for the intended intranasal delivery route. HPMC was prepared as a 1% w/v stock solution in Water for Injection (WFI) and added to the UF/DF2 filtrate at a volume calculated to yield a final drug substance HPMC concentration of 0.1% w/v. Following HPMC addition, the formulated drug substance was passed through a 0.45/0.2 µm bioburden reduction filter and dispensed into 1 L single-use bioprocess bags (Sartorius Flexboy), which were stored at 2 - 8 °C as the final IPD-52520 drug substance. The choice of HPMC concentration was based on precedent from intranasal protein formulations and its established compatibility with *E. coli*-derived recombinant proteins; formal excipient qualification was conducted as part of the formulation development program (to be published separately).

### 4.3 Endotoxin Control and Process Economics

The dual-stage endotoxin control strategy, polyethyleneimine (PEI) treatment during solubilization followed by downstream polishing, achieves robust endotoxin clearance to levels consistent with therapeutic requirements while maintaining high protein recovery. The overall process, comprising guanidine-based solubilization, tangential flow filtration (TFF)-mediated refolding, single-step anion exchange chromatography, ultrafiltration/diafiltration (UF/DF), and endotoxin removal, is intentionally streamlined. This design minimizes infrastructure complexity, supports efficient technology transfer, and is well-suited for scalable manufacturing.

The process achieves volumetric yields of approximately 2 g/L at >90% purity. This performance aligns with reported production ranges for recombinant proteins expressed in *E. coli*, where inclusion body-based processes and high-expression systems can drive product accumulation to 10–50% of total cellular protein and enable gram-per-liter titers under optimized fermentation conditions [27,28]. Accordingly, the yield obtained here falls within the expected high-performance range for small recombinant proteins produced via microbial systems.

## 5. Candidate Downselection

### 5.1 Rationale for Lead Candidate Selection

Process development described in Sections 5.2 to 5.4 was conducted in parallel on both the Trimer (IPD-52520; C389-AHB2v1-2GS-SB175, 17 kDa) and Daisy (AHB2v2_12PAS_LCB3v2.2_12PAS_LCB1v2.2, 25 kDa) miniprotein candidates to establish a comprehensive comparative dataset for informed candidate selection. While both molecules demonstrated acceptable manufacturability on the optimized *E. coli* BL21(DE3) platform, the Trimer candidate was selected as the lead for advancement into cGMP clinical trial material (CTM) manufacturing based on a combined assessment of preclinical efficacy, biophysical properties, and process performance.

### 5.2 Preclinical and Biophysical Basis for Downselection

Downselection criteria encompassed three domains: binding dissociation kinetics, neutralization breadth across SARS-CoV-2 variants of concern (VOCs), and expression and solubility. IPD-52520 (Trimer) was evaluated against these criteria using in vitro neutralization assays spanning multiple VOCs, a murine SARS-CoV-2 challenge model, and a pilot-scale manufacturability assessment covering purified yield and drug substance stability. Based on this integrated evaluation, IPD-52520 was selected to advance into cGMP development.

In preclinical studies, IPD-52520 demonstrated superior in vivo neutralization potency compared with Daisy. This advantage stems from its homotrimeric architecture, which presents three RBD-binding domains in a multivalent configuration, thereby enhancing avidity-driven target engagement—a critical determinant of therapeutic efficacy against rapidly evolving SARS-CoV-2 variants. IPD-52520 also exhibited favorable pharmacokinetic properties relative to Daisy, consistent with its smaller molecular footprint (17 kDa) and suitability for the intended intranasal delivery route.

From a manufacturability standpoint, IPD-52520 achieved higher volumetric yields at pilot scale: 2.5–2.9 g/L at 93.1–95.8% purity—compared with Daisy’s 1.7–2.0 g/L at the equivalent 10 L scale (Table 6). Beyond titer, IPD-52520 showed more consistent behavior across the inclusion-body solubilization and TFF refolding unit operations and produced a well-defined, reproducible elution profile on the DEAE-FF AEX platform, enabling reliable impurity clearance. Together, superior preclinical performance, higher manufacturing titers, and greater process robustness established an unambiguous advantage for IPD-52520 as the lead candidate.

### 5.3 Designation and Transition to cGMP Manufacturing

Based on these integrated assessments, the Trimer miniprotein was designated IPD-52520 and advanced as the sole lead candidate into cGMP manufacturing. All subsequent 50 L scale-up runs, clinical material production, analytical characterization, and stability studies described in Sections 6–8 pertain exclusively to IPD-52520 (Trimer). The process parameters and operating ranges established during the dual-candidate development phase (Sections 2 - 4) were directly translated to the cGMP process, with scale-dependent adjustments limited to equipment configuration and batch-size-related volumetric scaling. No fundamental changes to the unit operations or their sequence were required, confirming that the process platform developed during the parallel evaluation phase was robust and transferable.

## 6. cGMP Scale-Up and Clinical Material Production

### 6.1 Upstream Manufacturing Process

IPD-52520 drug substance (Batches: non-GMP, GMP01, GMP02, GMP06, GMP07, GMP09) was produced at 50 L scale under cGMP conditions in the same facility used for non-GMP process development. Batch non-GMP was a lot intended for non-clinical and reference standard studies. The remaining batches were manufactured under GMP conditions and designated for Phase 1 clinical studies.

The WCB thaw, first seed (500 mL flask, SY medium, 37 °C / 200 rpm, OD₆₀₀ ≥4.0), and second seed (4 × 1 L flasks, OD₆₀₀ ≥4.0) expansions preceded inoculation of a 75 L fermenter containing 25 L of SY medium (sterilized at 121.5 °C for 30 min). Fermentation was conducted at 37 °C, pH 6.95 (3 M NaOH), and DO 30% (cascade control, 200 - 500 rpm, 0 - 40 SLPM O₂ sparging). IPTG induction (50 mL of 0.5 M) was triggered at OD₆₀₀ ≥60, with harvest 6 h post-induction. Cell pellets (≥3 kg) were recovered by centrifugation (12,000 x g, 15 min, 4 °C) and stored at ≤ −70 °C.

### 6.2 Downstream Manufacturing Process

Frozen cell pellets were solubilized in DPBS containing 6 M guanidine-HCl and 0.05% PEI (5 L/kg pellet, 200 rpm, RT, 4 h), then clarified by centrifugation (17,000 x g, 60 min, 4 °C). Refolding was performed by UF/DF (30 kDa MWCO, ≤1 bar TMP) with five sequential dilution–concentration cycles against DPBS until permeate conductivity reached ≤20.0 mS/cm. Secondary clarification (15,000 x g, 30 min; 0.5/0.2 µm filtration) and a second PEI precipitation (0.03% v/v, ≥2 h) further reduced endotoxin. AEX chromatography (DEAE Sepharose FF; linear NaCl gradient 0–1 M in DPBS) was followed by HIC polishing (Phenyl Sepharose 6 FF; ammonium sulfate gradient 1–0 M). Final UF/DF concentrated the product to 66.0 g/L in DPBS, and 0.1% HPMC was added as a stabilizer. The drug substance was filtered through 0.45/0.2 µm filters and stored in 1 L bioprocess bags at 2–8 °C.

### 6.3 Batch Performance and Comparability

Eight independent batches were manufactured, with the first three batches supporting GLP toxicology studies and the remaining batches supplying Phase 1 clinical needs. Critical quality attributes: identity, purity, impurity profiles, protein concentration, and overall yield, showed excellent batch-to-batch comparability with no scale-dependent trends (**Table 8**, **Table 9**). These data confirm that the *E. coli* expression and purification platform delivers reliable, consistent production at clinical scale.

**Table 7.** Summary of candidate comparison supporting Trimer downselection.

| **Attribute** | **Trimer (IPD-52520)** | **Daisy (IPD-52521)** |
| --- | --- | --- |
| Molecular weight | 17 kDa | 25 kDa |
| Architecture | Homotrimer (multivalent) | Tandem fusion |
| Pilot-scale yield (10 L) | 2.4–2.9 mg/L | 1.7–1.9 mg/L |
| Pilot-scale purity | 93.1–95.8% | 94.4–98.7% |
| In vivo neutralization | Superior | Acceptable |
| PK / Biodistribution | Favorable (smaller size) | Adequate |
| Process consistency | High | Moderate (variable lysis) |
| Advancement decision | Selected for Clinical development | Discontinued |

**Table 8.**
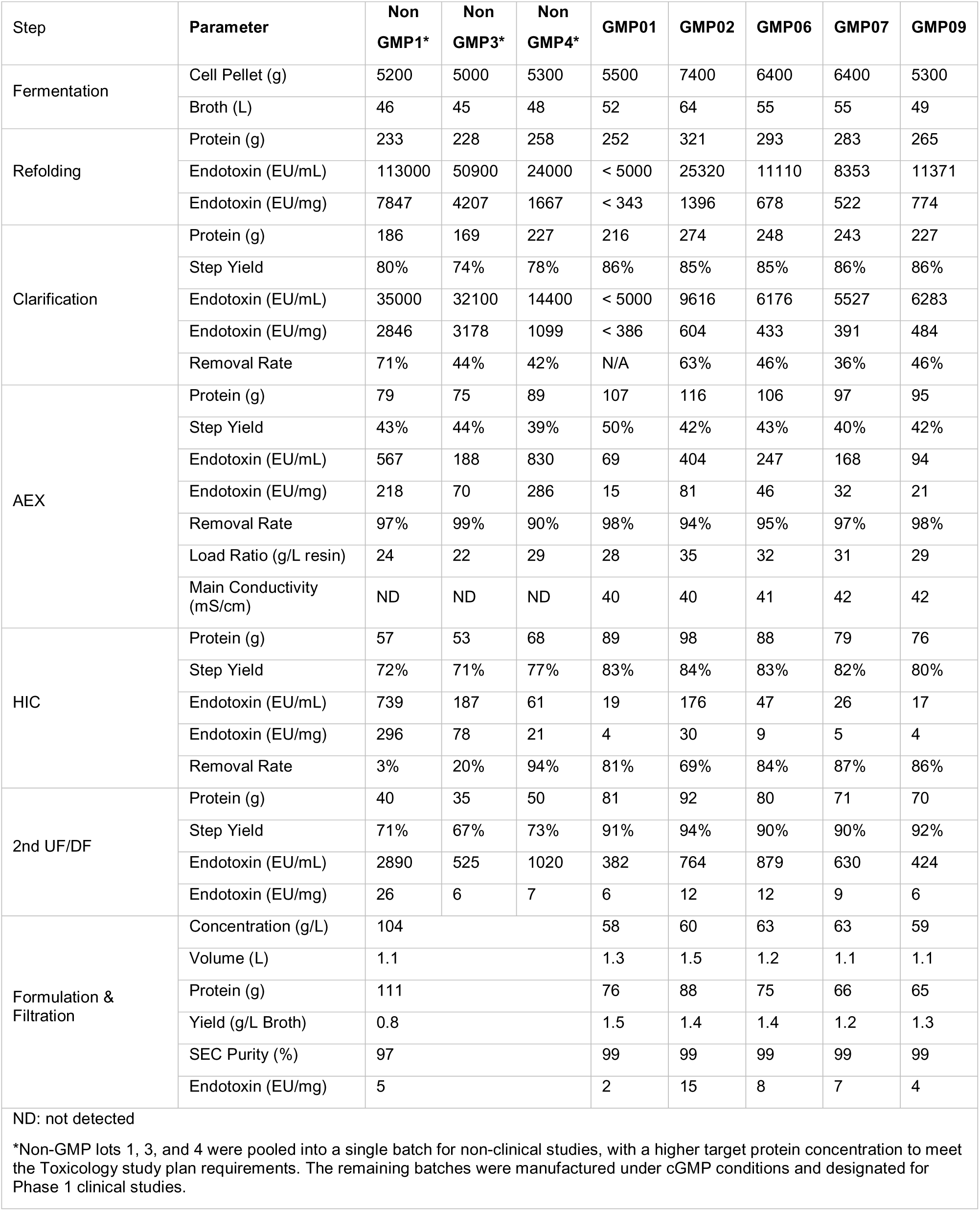
Process mass balance and step yield for IPD-52520 downstream manufacturing. Results are shown for each lot individually. Non-GMP lots 1, 3, and 4 were co-processed as a single pooled drug substance campaign at the formulation stage (shown as pooled values). Endotoxin removal rates are relative to the preceding unit operation pool. ND: not detected. N/A: not applicable (pooled lot).

**Table 9.**
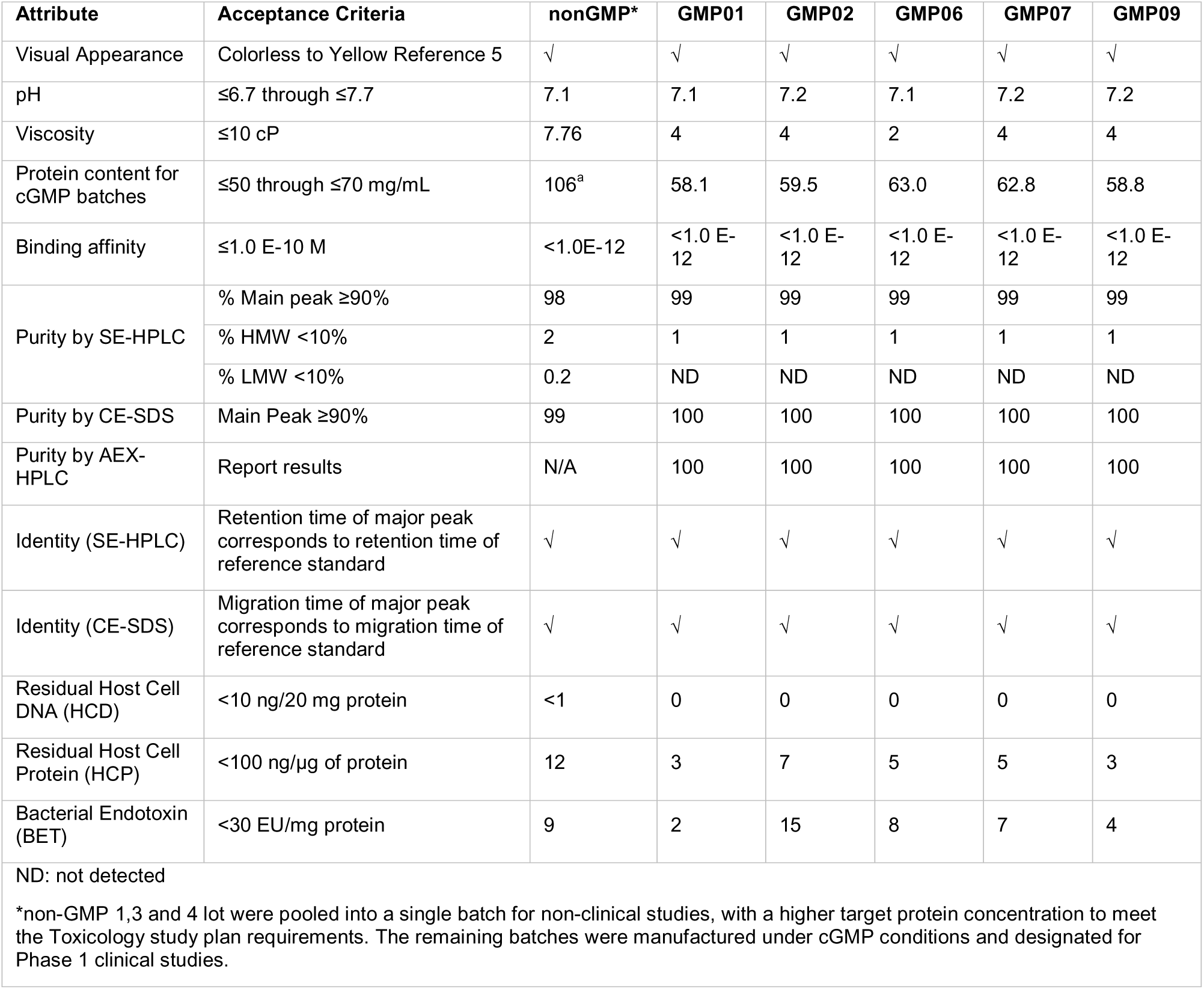
Summary of quality attributes across full-scale (50L) batches.

#### 6.3.1 Process Mass Balance and Step Yield

The downstream process mass balance, covering all unit operations from the solubilized cell pellet pool to the final drug substance, is summarized in Table 8. Results are presented separately for each of the three non-GMP lots (Lots 1, 3, and 4) and for each of the five cGMP batches. For the drug substance formulation step, the non-GMP lots were co-processed as a single pooled downstream campaign and are reported accordingly.

##### Overall process yield

Across all five cGMP batches, the optimized downstream process delivered final drug substance yields of 1.2–1.5 g/L of fermentation broth, with ≥99% SE-HPLC main-peak purity and endotoxin levels of 2–15 EU/mg, well within the proposed acceptance limit of 30 EU/mg. The non-GMP pooled campaign achieved a lower volumetric yield of 0.8 g/L of broth, reflecting the inclusion of earlier-stage development lots with less-optimized process parameters. It was intentionally concentrated to 104 g/L drug substance to meet the non-clinical toxicology study dosing plan.

##### AEX chromatography

The AEX capture step (DEAE Sepharose FF) consistently delivered the lowest individual step yield in the process, recovering 39–50% of the loaded protein across all lots. This step yield reflects the deliberate use of stringent wash and gradient elution conditions required to meet purity and endotoxin clearance objectives, as AEX provided the single largest endotoxin reduction across the process (90–99% removal per step). Despite the moderate step yield, resin loading ratios were well controlled (22–35 g protein per liter of resin), confirming that yield losses at this step are attributable to process selectivity rather than capacity overloading.

##### Endotoxin clearance cascade

Endotoxin levels entering the downstream process from the refolded pool were substantial, ranging from 522–7,847 EU/mg across all lots, consistent with the high endotoxin burden typical of *E. coli* inclusion-body processing. The process employed an orthogonal, multi-step clearance strategy; PEI precipitation during solubilization, further reduction at the clarification step (36–71% removal), major clearance at AEX (90 - 99% removal), and additional polishing at HIC, to achieve a final drug substance endotoxin level of 2–15 EU/mg: a total process reduction of greater than 3 log_10_ relative to the refolded pool. Endotoxin profiles at the HIC step showed greater variability in the non-GMP lots (3 - 94% removal) compared with the cGMP batches (69–87% removal), a direct consequence of the Tergitol AEX wash concentration optimization described in Section 4.2.1. The transition to 3% Tergitol in the cGMP process delivered substantially more consistent HIC endotoxin clearance.

##### Non-GMP to cGMP process improvement

Step yields improved markedly from the non-GMP development lots to the cGMP batches across multiple unit operations. Clarification step yield increased from 74–80% (non-GMP) to 85–86% (cGMP); HIC step yield from 71–77% to 80–84%; and, most substantially, the second UF/DF step yield from 67–73% to 90–94%. These improvements reflect process refinements implemented between the non-GMP confirmation runs and cGMP manufacture, including optimized centrifugation parameters, membrane selection, and TMP control during the UF/DF2 concentration step. The five cGMP batches demonstrated highly consistent step yields across all unit operations, confirming process robustness at clinical manufacturing scale.

## 7. Material Characterization

IPD-52520 (drug substance from a non-GMP lot) was characterized using the analytical and biochemical methods summarized in Table 10. Detailed procedures and results for each technique are provided in the subsections below.

**Table 10.** Characterization methods and results for IPD-52520.

| Attribute | Analytical Method | Result |
| --- | --- | --- |
| Molecular weight, polydispersity, and hydrodynamic radius | SEC-MALS / QLS / dRI | Main peak ~46 kDa; pDI = 1.005; Rh = 5.6 nm |
| Amino acid composition | Acid hydrolysis / UPLC | 139 residues confirmed (Trp excluded); ≥91% recovery for all residues |
| Primary structure / intact mass | LysC/Trypsin LC-MS/MS | Main observed mass 16,602 Da (theoretical 16,601.6 Da) |
| Thermal stability | Differential scanning calorimetry (DSC) | Single asymmetric unfolding peak; dominant T<sub>m</sub> = 73.4 °C |
| Secondary and tertiary structure | Circular dichroism (far-UV and near-UV) | α-Helix 85.4%; β-Sheet 0.0%; β-turn 2.6%; Other 12.0%; defined aromatic signatures by near-UV CD |

### 7.1 Molecular Weight, Polydispersity, and Hydrodynamic Radius

The molecular weight (MW), polydispersity, and hydrodynamic radius of IPD-52520 were determined by size-exclusion chromatography coupled with multi-angle light scattering (SEC-MALS), quasi-elastic light scattering (QLS), and differential refractive index (dRI) detection. Samples were chromatographed on a Tosoh TSKgel G3000SWXL column (5 µm, 7.8 mm × 300 mm) using an Agilent 1200 HPLC system equipped with an inline 0.1 µm filter. Analytes eluting from the SEC column passed sequentially through four detectors: UV absorbance (280 nm), MALS for MW determination, QLS for hydrodynamic radius, and dRI for absolute MW and polydispersity calculations. A refractive index increment of dn/dc = 0.185 mL/g was used. All analyses were performed in duplicate.

The SEC-MALS chromatogram is shown in **Figure 9**. Molecular weight values are plotted across the eluting peak; edge inflections are attributable to reduced signal-to-noise at the peak flanks. The weighted-average MW of the main peak was approximately 46 kDa, within the expected range for the predicted molecular mass of the IPD-52520 assembly. The polydispersity index (pDI) of 1.005, derived from the number-average MW, confirms a homogeneous, essentially monodisperse population. The QLS-derived hydrodynamic radius of 5.6 nm for the main peak is consistent with the expected hydrodynamic behavior of this molecular assembly in solution.

**Figure 9.**
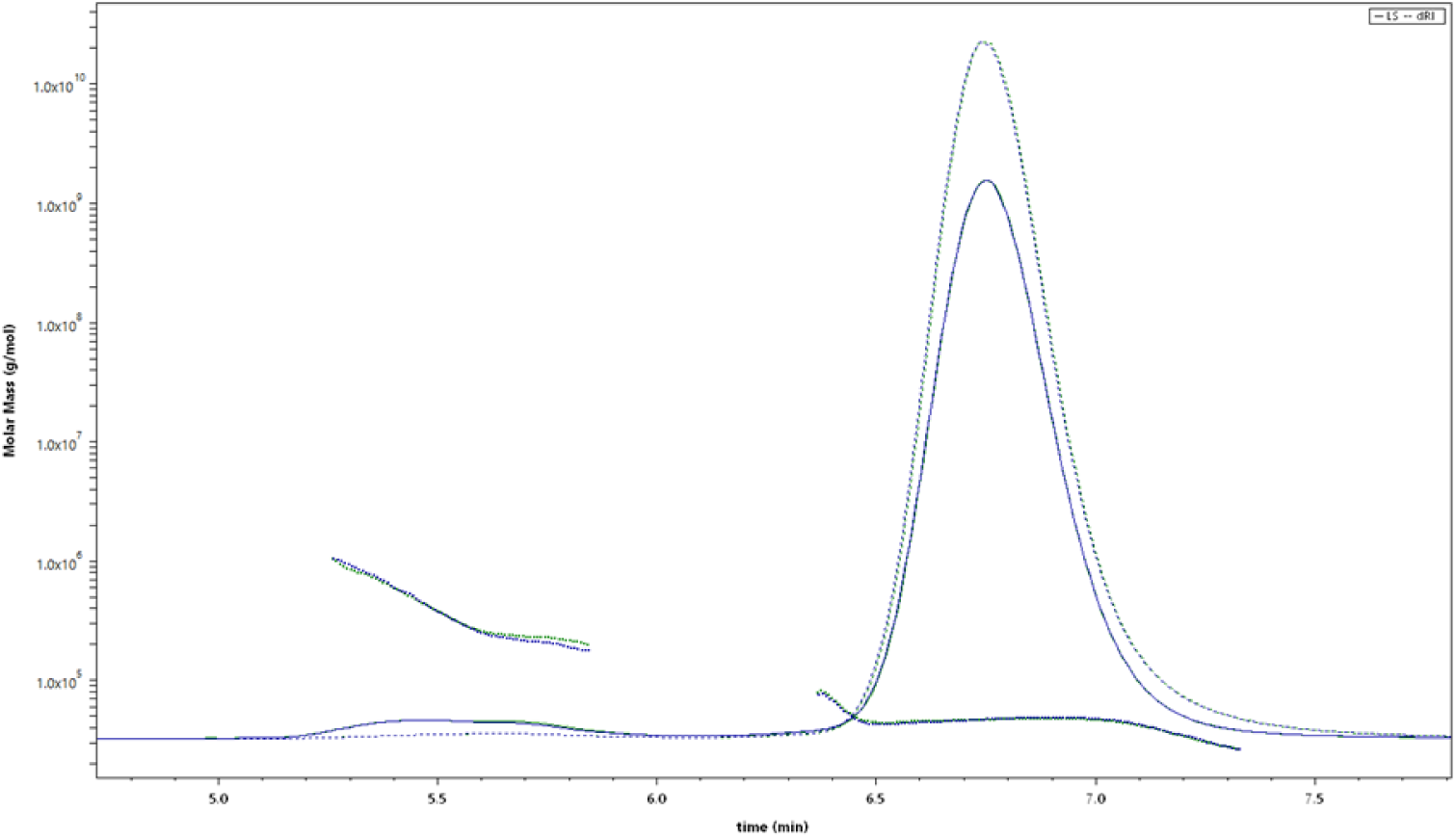
SEC-MALS analysis of IPD-52520. Representative chromatogram showing 90° light scattering (solid line) and differential refractive index (dotted line), with molecular weight plotted across the main peak (scatter, right axis, g/mol). Weighted-average MW ≈ 46 kDa; pDI = 1.005; Rh = 5.6 nm. Drug substance material from a non-GMP lot; analysis performed in duplicate.

### 7.2 Primary Structure Confirmation

#### 7.2.1 Amino Acid Composition

Amino acid composition was determined by acid hydrolysis followed by UPLC analysis. Samples were hydrolyzed with 4% (v/v) phenol in 6 N HCl under vacuum for 22 h (Eldec Pico Tag Workstation) and then analyzed on an Acquity UPLC BEH C18 column (1.7 µm, 2.1 × 100 mm) with UV detection at 260 nm. Analyses were performed in triplicate. The analysis confirmed 139 amino acid residues, consistent with the theoretical sequence (tryptophan is destroyed under acid hydrolysis conditions and was excluded). Recovery was 98–103% for the six most abundant residues (Arg, Asx, Glx, Ala, Lys, and Leu), 91–98% for residues present in four or more copies, and within the accepted analytical range for residues present in one to three copies. These results confirm that the amino acid composition of IPD-52520 is consistent with the designed sequence.

#### 7.2.2 Peptide Mapping and Intact Mass Analysis by LC-MS/MS

Primary structure was confirmed by LC-MS/MS peptide mapping with dual-enzyme digestion and by intact mass spectrometry. For peptide mapping, 200 µg of IPD-52520 was reconstituted in 6 M guanidine-HCl / 250 mM Tris, pH 7.5, reduced with 50 mg/mL dithiothreitol (60 °C, 20 min), and alkylated with 0.5 M iodoacetamide (RT, 30 min, protected from light). Samples were diluted into 50 mM Tris, pH 7.5, and digested separately with LysC (0.5 mg/mL, 25 °C, 18 h) and trypsin (0.5 mg/mL, 25 °C, 18 h). Reactions were quenched with 20% formic acid. Resulting peptides were resolved on a Waters Acquity Peptide CSH C18 column (1.7 µm, 2.1 × 150 mm, Waters P/N 186006938) using an acetonitrile / 0.085% formic acid gradient. MS and MS/MS spectra were acquired on a Waters Xevo G2-XS QTof instrument and processed with Waters UNIFI software. Full sequence coverage was obtained.

The deconvoluted intact mass spectrum is shown in **Figure 10**. The principal observed mass of 16,602 Da is in excellent agreement with the theoretical monoisotopic mass of 16,601.6 Da (Δ < 1 Da), confirming the correct primary structure and the absence of significant post-translational modifications. Minor mass variants detected at low relative abundance are summarized in Table 10; these range from 16,630 to 16,942 Da and are consistent with chemical modifications commonly observed in *E. coli*-derived recombinant proteins.

**Figure 10.**
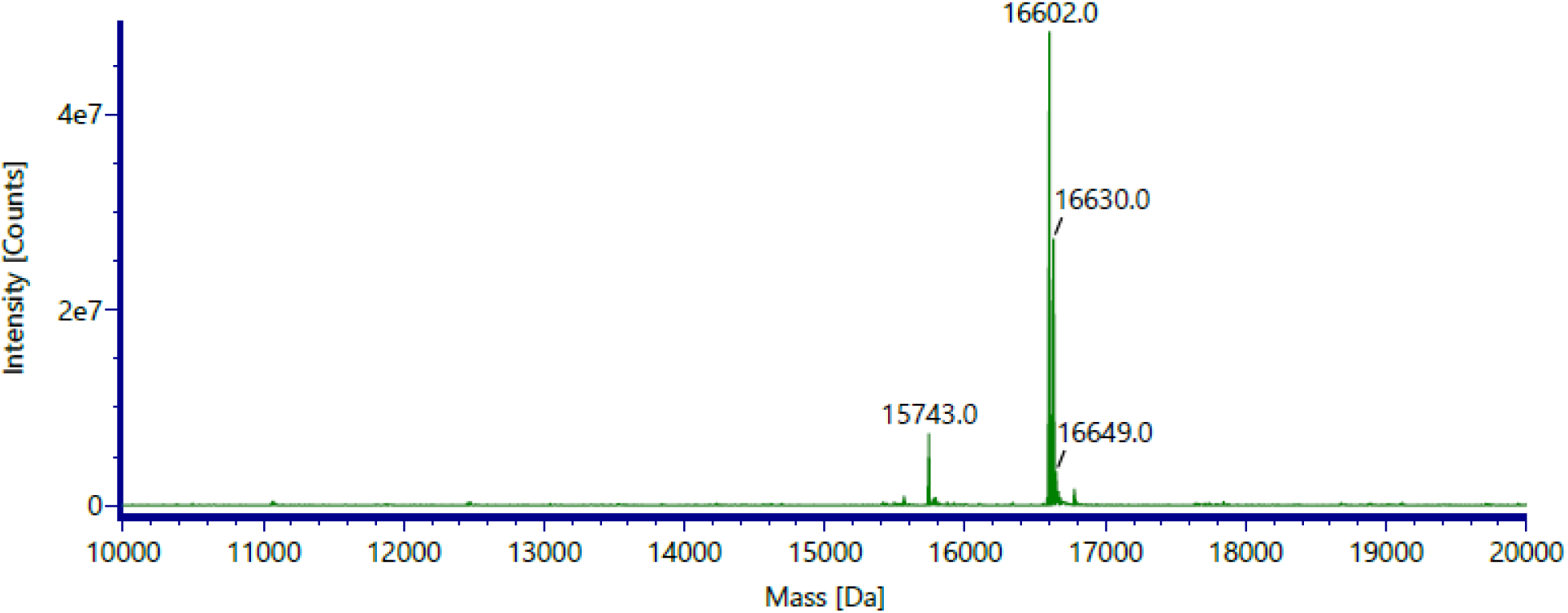
Deconvoluted intact mass spectrum of IPD-52520. The dominant species at 16,602 Da corresponds to the native, unmodified protein chain. Minor mass variants detected at low relative abundance are annotated; their observed masses are listed in Table 11.

**Table 11.** Intact mass analysis of IPD-52520. All minor variants were detected at low relative abundance.

| Expected Mass (Da) | Observed Mass (Da) | Notes |
| --- | --- | --- |
| 16,601.6 | **16,602** | Native — principal species |
| — | 16,630 | Minor species |
| — | 16,643 | Minor species |
| — | 16,648 | Minor species |
| — | 16,677 | Minor species |
| — | 16,779 | Minor species |
| — | 16,942 | Minor species |

### 7.3 Thermal Stability by Differential Scanning Calorimetry

Thermal unfolding behavior was characterized by DSC. IPD-52520 was diluted to 4.75 mg/mL in formulation buffer (DPBS + 0.1% HPMC w/v). Heat-capacity data were corrected relative to the formulation buffer baseline and converted to molar heat capacity (kJ/mol·K) using the measured protein concentration and theoretical molecular weight. The resulting thermogram is shown in Figure 11.

**Figure 11.**
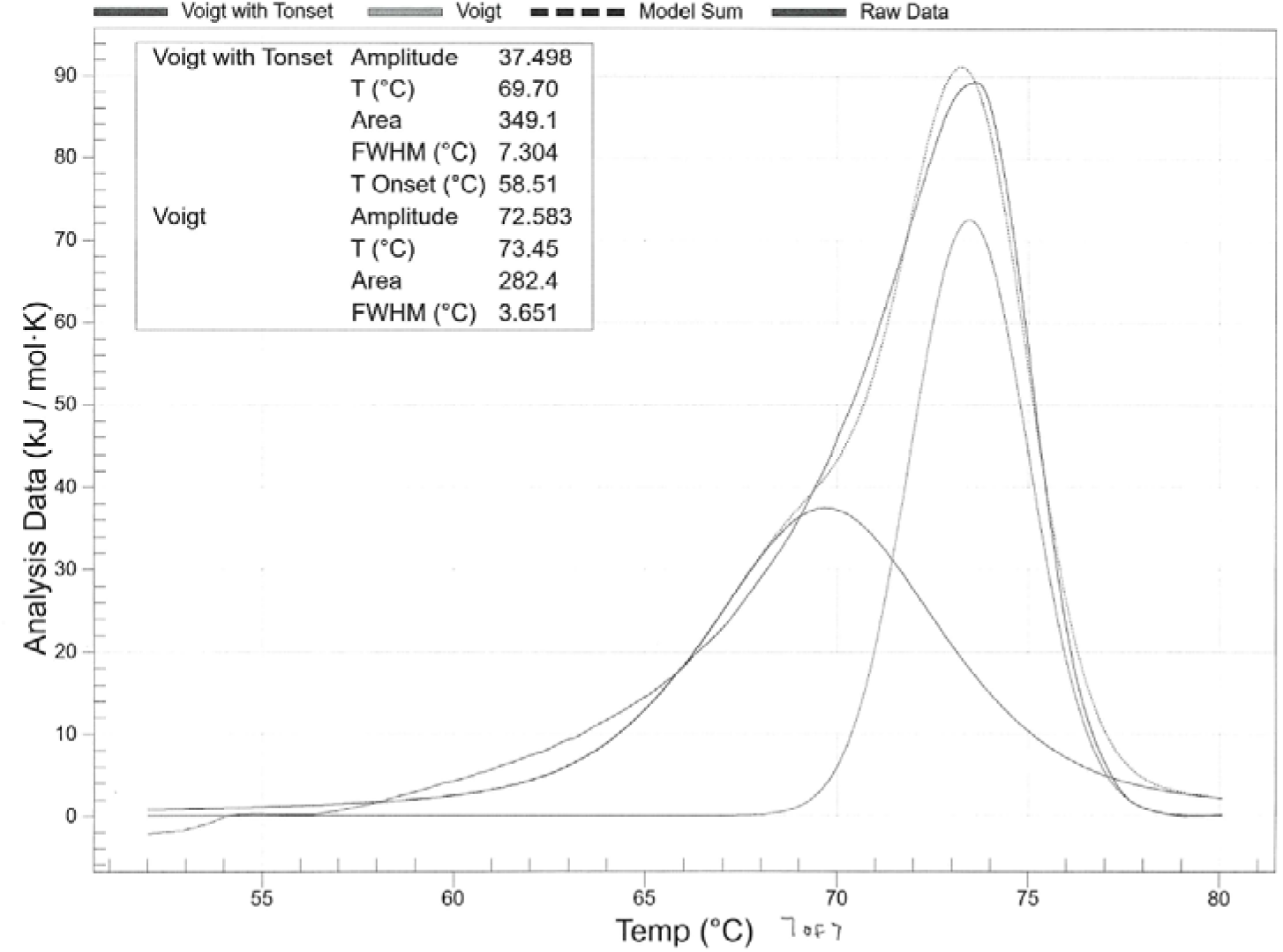
DSC thermogram of IPD-52520. Baseline-corrected molar heat capacity data (black solid line), two-state scaled model fit (dotted line), individual Voigt model fits (dashed lines), and their sum (colored solid line). IPD-52520 at 4.75 mg/mL in DPBS + 0.1% HPMC formulation buffer; scan rate 1 °C/min.

A single endothermic peak with pronounced asymmetry was observed. A two-state scaled model did not fit the experimental data well, indicating that IPD-52520 does not unfold through a simple cooperative transition and that one or more intermediate states are populated during denaturation. Two Voigt models were applied sequentially; their sum closely reproduced the experimental thermogram and provided a substantially better fit than the two-state scaled model. The dominant unfolding transition midpoint of 73.4 °C reflects the primary cooperative unfolding event and represents the operational thermal stability of the intact assembly. A minor pre-transition feature near a 54 °C onset is attributed to partial unfolding of a less stable structural element preceding global denaturation. A T_m_ of 73.4 °C demonstrates excellent thermal stability for a miniprotein therapeutic and provides a wide margin above the intended 2–8 °C storage condition.

### 7.4 Secondary and Tertiary Structure by Circular Dichroism

The secondary and tertiary structures of IPD-52520 were evaluated by circular dichroism (CD) spectroscopy. Far-UV CD (190–260 nm) was measured at 0.25 mg/mL using a 0.1 cm pathlength cell; near-UV CD (240–350 nm) was measured at 1.0 mg/mL using a 1.0 cm pathlength cell. All spectra were corrected against a matched blank buffer. Data are expressed as Mean Residue Ellipticity (MRE, deg·cm^2^·dmol^−1^). Quantitative secondary structure deconvolution was performed using Principal Component Regression (PCR) analysis; results are summarized in **Table 12**.

**Table 12.** Secondary structure composition of IPD-52520 by far-UV CD. Derived by Principal Component Regression (PCR) deconvolution.

| Sample | α-Helix (%) | β-Sheet (%) | Turn (%) | Other (%) |
| --- | --- | --- | --- | --- |
| IPD-52520 | 85.4 | 0.0 | 2.6 | 12.0 |

The far-UV CD spectrum (Figure 12) shows the canonical signature of a predominantly α-helical protein: a positive band at 191.7 nm (MRE = +73,812 deg·cm^2^·dmol^−1^) and negative minima at 208.5 nm (MRE = −32,539 deg·cm^2^·dmol^−1^) and 221.0 nm (MRE = −3,122 deg·cm^2^·dmo^l−1^), corresponding to the π→π* and n→π* amide transitions. PCR deconvolution confirmed 85.4% α-helical, 0.0% β-sheet, 2.6% turn, and 12.0% other, fully consistent with the designed all-helical architecture.

**Figure 12.**
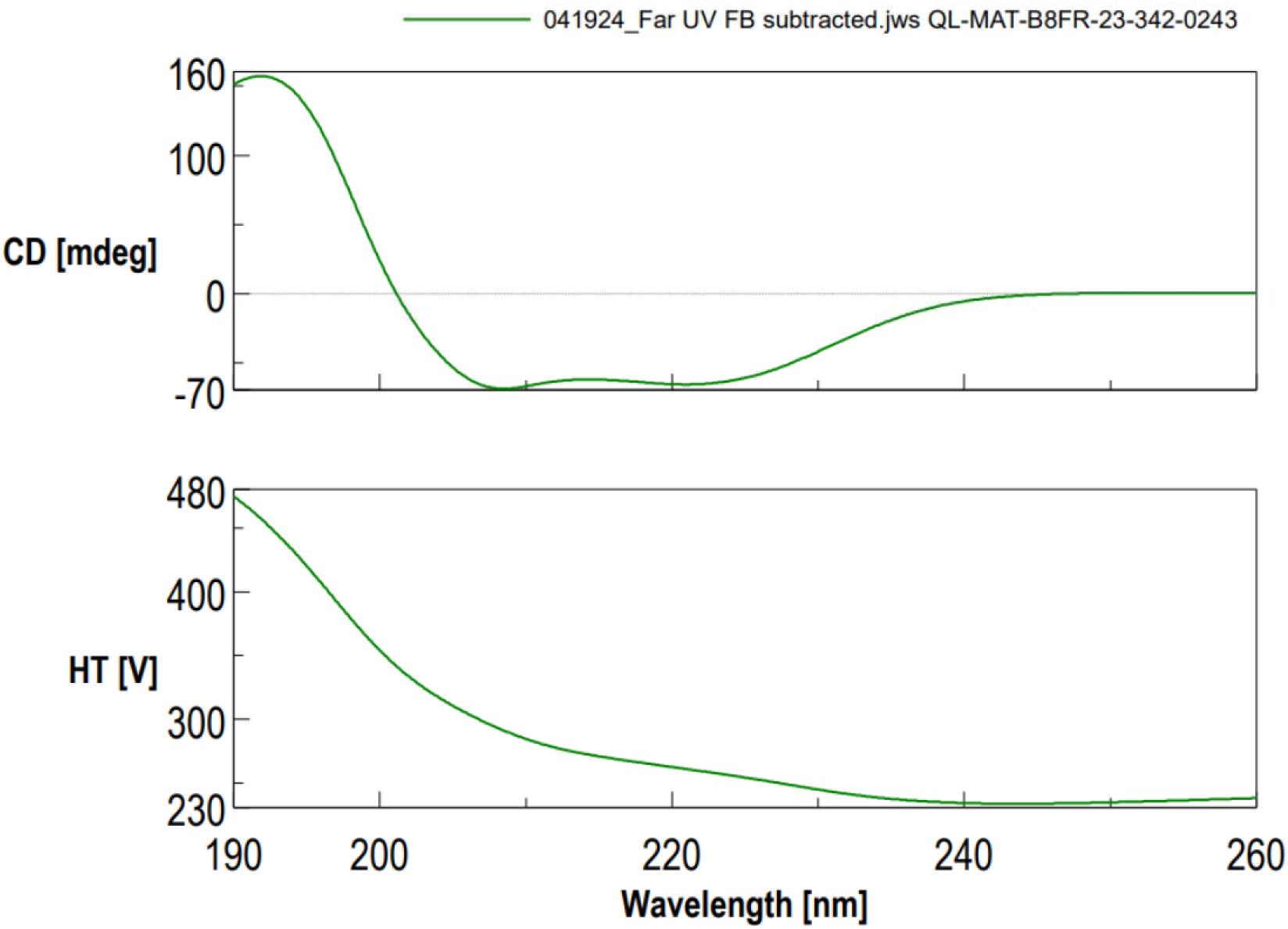
Far-UV CD MRE spectrum of IPD-52520. Mean Residue Ellipticity as a function of wavelength (190–260 nm), recorded at 0.25 mg/mL in a 0.1 cm pathlength cell and corrected against a matched blank buffer. A positive band at 191.7 nm and negative minima at 208.5 nm and 221.0 nm confirm a predominantly α-helical secondary structure.

The near-UV CD spectrum (240–350 nm) showed contributions from phenylalanine (250–270 nm), tyrosine (270–290 nm), and tryptophan (280–300 nm) side chains, resolved into six distinct features. The defined peak pattern indicates that aromatic residues occupy specific, restricted environments within the folded structure, consistent with their designed burial within the hydrophobic core. The modest overall signal amplitude is characteristic of small, entirely α-helical proteins lacking disulfide bonds and does not indicate the absence of tertiary structure.

### 7.5 Characterization Summary and Conclusions

The combined orthogonal analytical data provide a thorough and consistent characterization of IPD-52520. The identity and primary structure are verified: the 139-residue amino acid sequence matches the theoretical one, and the main intact mass of 16,602 Da aligns with the calculated monoisotopic mass of 16,601.6 Da within 1 Da. Minor mass variants observed at low levels are typical of chemical modifications common in *E. coli*-derived recombinant proteins and do not affect product identity. The higher-order structure is well defined and aligns with the intended design. Far-UV CD indicates 85.4% α-helical secondary structure, consistent with the computational model. Near-UV CD shows aromatic residues are in specific, well-defined environments within the folded core. DSC reveals a primary unfolding transition at *T_m_* = 73.4 °C, providing a significant thermal margin above the 2–8 °C storage temperature. Colloidal properties are favorable: SEC-MALS confirms a monodisperse population (pDI = 1.005) with an average molecular weight of about 46 kDa and a hydrodynamic radius of 5.6 nm. Overall, these findings demonstrate that the *E. coli* expression and purification process produces IPD-52520 with the structural integrity, uniformity, and biophysical characteristics necessary for clinical development.

## 8. Stability

A comprehensive stability program evaluated IPD-52520 drug substance under real-time (5 ± 3 °C) and accelerated (25 ± 2 °C / 60 ± 5% RH) conditions.

### 8.1 Real-Time Stability (5 ± 3 °C)

Lot #GMP07 maintained robust physicochemical and functional stability over 12 months (Table 13). Visual appearance, pH (7.1–7.2), protein concentration (62.3–64.6 mg/mL), and viscosity (4 cP) remained stable throughout. SE-HPLC purity remained at 98–99%, with HMW species at 1–2% and no detectable LMW fragments. AEX-HPLC main peak purity was 99.99–100%. BLI binding affinity remained below the quantitation threshold of 1.0 × 10⁻¹² M at all time points. Residual DNA, HCP, endotoxin, and bioburden were consistently within specifications.

**Table 13.** Lot #GMP07 stability at 5 ± 3 °C (12-month data; study ongoing).

| Test Method | Acceptance criteria | Time Point (Months) |  |  |  |  |  |
| --- | --- | --- | --- | --- | --- | --- | --- |
|  |  | 0 | 1 | 3 | 6 | 9 | 12 |
| Visual Appearance | Colorless to Yellow (Ref. 5) | √ | √ | √ | √ | √ | √ |
| pH | $6.7 \leq \text{pH} \leq 7.7$ | 7.2 | 7.1 | 7.1 | 7.1 | 7.1 | 7.1 |
| Protein Content | 50.0–70.0 mg/mL | 62.8 | 63.1 | 62.3 | 63.4 | 64.6 | 62.5 |
| Binding Affinity (BLI) | $\leq 1.0\text{E-}10$ M | $< 1.0\text{E-}12$ M | $< 1.0\text{E-}12$ M | $< 1.0\text{E-}12$ M | $< 1.0\text{E-}12$ M | $< 1.0\text{E-}12$ M | $< 1.0\text{E-}12$ M |
| Viscosity | $\leq 10$ cP | 4 cP | NT | NT | NT | NT | 4 cP |
| Purity SE-HPLC | % Main Peak $\geq 90\%$ | 99% | 98% | 98% | 98% | 98% | 99% |
| | % HMW $\leq 10\%$ | 1% | 2% | 2% | 2% | 2% | 1% |
| | % LMW $\leq 10\%$ | NT | NT | NT | NT | NT | NT |
| Purity CE-SDS (NR) | Major Peak $\geq 90\%$ | 100% | NT | NT | NT | NT | 100% |
| Purity AEX-HPLC | % Main Peak- Report Results | 100% | 100% | 100% | 100% | 100% | 99.99% |
| Identity SE-HPLC | RT vs. Reference-Pass | √ | NT | NT | NT | NT | √ |
| Identity CE-SDS (NR) | MT vs. Reference-Pass | √ | NT | NT | NT | NT | √ |
| Residual DNA | $< 10$ ng/20 mg | 0 | NT | NT | NT | NT | 0 |
| Residual HCP | $< 100$ ng/μg | 5 ng/μg | NT | NT | NT | NT | 6 ng/μg |
| Bacterial Endotoxin | $< 30$ EU/mg | 7 EU/mg | NT | NT | NT | NT | 3 EU/mg |
| Bioburden | TAMC $\leq 10^2$ CFU/mL | 0 CFU/mL | NT | NT | NT | NT | 0 |
| | TYMC $\leq 10^1$ CFU/mL | 0 CFU/mL | NT | NT | NT | NT | 0 |

NT = Not tested at this time point.
√ - Pass

### 8.2 Accelerated Stability (25 ± 2 °C / 60 ± 5% RH)

Most attributes remained within specification through 6 months (Table 14). Protein concentration increased from 62.8 to 72.3 mg/mL, exceeding the upper limit at 6 months, most likely due to evaporative volume loss from the container closure system under thermal stress rather than a protein-intrinsic pathway, as no corresponding degradation signals were observed. A transient HMW increase at 3 months (4%, reverting to 1% at 6 months) may reflect assay variability rather than a true aggregation event. AEX-HPLC purity shifted modestly (100% to 99.79%), consistent with minor charge-variant accumulation under stress.

**Table 14.** Accelerated stability at 25 ± 2 °C / 60 ± 5% RH.

| Test Method | Acceptance Criteria | Time Point (Months) |  |  |  |
| --- | --- | --- | --- | --- | --- |
|  |  | 0 (lot release) | 1 | 3 | 6 |
| Visual Appearance | Colorless to Yellow (Ref. 5) | √ | √ | √ | √ |
| pH | 6.7 ≤ pH ≤ 7.7 | 7.2 | 7.1 | 7.1 | 7.1 |
| Protein Content | 50.0–70.0 mg/mL | 62.8 mg/mL | 64.1 mg/mL | 66.2 mg/mL | 72.3 mg/mL* |
| Binding Affinity (BLI) | ≤ 1.0E-10 M | < 1.0E-12 M | < 1.0E-12 M | < 1.0E-12 M | < 1.0E-12 M |
| Viscosity | ≤ 10 cP | 4 cP | 4 cP | 4 cP | 4 cP |
| Purity SE-HPLC | % Main Peak ≥ 90% | 99% | 99% | 96% | 99% |
|  | % HMW ≤ 10% | 1% | 2% | 4% | 1% |
|  | % LMW ≤ 10% | NT | NT | NT | NT |
| Purity CE-SDS (NR) | Major Peak ≥ 90% | 100% | 100% | 100% | 100% |
| Purity AEX-HPLC | % Main Peak<br>Report Results | 100% | 100% | 100% | 99.79% |
| Identity SE-HPLC | RT vs. Reference-Pass | √ | NT | NT | √ |
| Identity CE-SDS (NR) | MT vs. Reference-Pass | √ | NT | NT | √ |

NT = Not tested at this time point.
\*Protein content of 72.3 mg/mL at 6 months exceeds the upper specification limit of 70.0 mg/mL; attributed to evaporative volume loss under stress conditions rather than intrinsic protein instability.

### 8.3 Stability Conclusions

IPD-52520 drug substance is stable at 5 ± 3 °C for at least 12 months, with no significant changes in purity, charge variant profile, or biological activity under either real-time or accelerated conditions. The excess protein content at 6 months under accelerated stress is attributable to container closure performance rather than to product degradation. These results provide a sound scientific basis for the proposed 2-8 °C storage condition and support continued clinical use within the defined shelf life.

## 9. Conclusions

This work presents the end-to-end development, cGMP manufacturing, characterization, and stability evaluation of IPD-52520, a *de novo*-designed antiviral miniprotein targeting SARS-CoV-2. Key achievements include:

1. A systematic fermentation optimization yielding a robust *E. coli* BL21(DE3) process with simplified SY medium, pH-stat feeding, and a late-induction strategy that reduces process time while maximizing expression.
2. A streamlined recovery and purification platform; guanidine solubilization, TFF refolding, single-column AEX, UF/DF, and dual-stage endotoxin removal, delivering approximately 2 g/L at >90% purity with minimal facility requirements.
3. Successful cGMP scale-up to 50 L, with eight independent batches demonstrating excellent comparability, supporting nonclinical toxicology and Phase 1 clinical supply.
4. Comprehensive biophysical characterization confirming a well-folded, thermally stable (α-helical content 85.4%; Tm 73.4 °C), monodisperse miniprotein with an intact primary structure and picomolar target-binding affinity.
5. A stability profile supports storage at 2 - 8 °C for at least 12 months, and the stability studies are ongoing.

These results demonstrate the feasibility of translating computationally designed antiviral proteins into manufacturable biologics and establish a platform adaptable to rapid-response therapeutics against emerging viral threats. Future development may explore alternative HMW removal strategies, continuous processing, and expanded clinical-scale manufacturing.

## Author contributions

SK Bioscience: Study protocols, execution, data generation, analysis, and manuscript authoring

Sammaiah Pallerla: Program Technical Oversight, Manuscript authoring.

Ryan Swoyer: Material characterization and analytical execution oversight.

IPD: Designed the minibinders.

Daniel Criag, Sandip Patel: Technical support and manuscript review.

Rashmi Ravichandran, Lauren Carter- Process Development and manuscript review

Praveen Alamuri: Program management support

All co-authors reviewed the manuscript

## Acknowledgments

This work was supported by funding from the Gates Foundation. The authors gratefully acknowledge the guidance and support of Jacqueline Kirchner, Deputy Director, as well as the colleagues on the CMC and analytical teams at the Gates Foundation, for their valuable input throughout the execution of this program..

The authors also acknowledge IAVI leadership’s support in coordinating and advancing this work. In addition, the authors recognize the support of the Leadership teams at SK Bioscience, which enabled the development and manufacturing activities described in this study.

